# A selectivity filter in the EMC limits protein mislocalization to the ER

**DOI:** 10.1101/2022.11.29.518402

**Authors:** Tino Pleiner, Masami Hazu, Giovani Pinton Tomaleri, Vy Nguyen, Kurt Januszyk, Rebecca M. Voorhees

**Author notes:** Neomorph Inc., 5590 Morehouse Dr., San Diego, CA 92121, USA. Equal contribution (Tino Pleiner, Masami Hazu, Giovani Pinton Tomaleri). Correspondence (RMV).

## Abstract

Tail anchored proteins (TAs) play essential roles at both the ER and mitochondria, and their accurate localization is critical to proteostasis. Biophysical similarities lead to mistargeting of mitochondrial TAs to the ER, where they are delivered to the ER membrane protein complex (EMC). We showed that the EMC directly contributes to sorting fidelity of mitochondrial TAs and multipass substrates that contain positively charged soluble domains. Leveraging an improved structural model of the human EMC, we used mutagenesis and site-specific crosslinking to map the path of a TA from its cytosolic capture by methionine-rich loops to its membrane insertion through a hydrophilic vestibule. Positively charged residues at the entrance to the vestibule function as a selectivity filter that uses charge-repulsion to reject mitochondrial TAs. Substrate discrimination by the EMC provides a biochemical explanation for one role of charge in TA sorting and protects compartment identity by limiting protein misinsertion.

## INTRODUCTION

A hallmark of eukaryotic cells is their organization into subcellular compartments that spatially separate otherwise incompatible biochemical reactions. The evolution of compartmentalization enabled the increasingly complex cellular processes required for emergence of multicellular life. To carry-out distinct functions, each compartment must contain a unique and precisely defined set of proteins and metabolites.

Membrane proteins comprise ~20% of the human proteome (Krogh et al., 2001), and their localization is a primary determinant of organellar identity, underscoring the importance of their accurate sorting. Due to the presence of one or more hydrophobic transmembrane domains (TMDs), targeting and insertion of membrane proteins must be tightly regulated to prevent their aggregation in the aqueous cytosol. Canonical localization of membrane proteins to their two major destinations in the cell, the mitochondria and endoplasmic reticulum (ER), relies on cleavable targeting sequences that direct proteins to the correct organelle. Both the mitochondrial targeting sequence (MTS) and the ER-specific signal sequence are proteolytically removed upon arrival at their respective compartment, and thus have evolved principally to ensure accurate sorting without the need to serve a functional role in the mature protein.

However, given the functional and topological diversity of the membrane proteome, many nascent proteins cannot utilize these stereotypical biogenesis pathways. In these cases, membrane proteins instead rely on recognition of a TMD and its surrounding residues for accurate sorting (Rapoport et al., 2017; Guna and Hegde, 2018). These sequences must therefore play dual roles, experiencing evolutionary pressure to both function in the mature protein (i.e. insertion, folding, and assembly) and ensure accurate localization.

One important family of membrane proteins that rely on their TMD and its flanking residues for recognition, targeting, and insertion are tail-anchored proteins (TAs) (Kutay et al., 1993; Chio et al., 2017; Hegde and Keenan, 2011; Guna et al., 2022a). TAs are characterized by a single C-terminal TMD followed by a short soluble domain of up to 30-40 amino acids. Their globular N-termini are localized to the cytosol and are responsible for carrying out their diverse functions. Because of their topology, the TMD of a TA emerges from the exit tunnel of the ribosome only after translation termination, and they must be post-translationally targeted to the correct organelle. TAs are found on all cellular membranes and regulate essential processes such as neurotransmitter release via exocytosis (SNARE proteins), cholesterol synthesis at the ER (squalene synthase [SQS]), and the onset of apoptosis at mitochondria (BCL-2, Bak). Given their biophysical diversity and the limited information for targeting, how TAs are accurately sorted between compartments has been a long-standing open question in the field.

TA localization is thought to be primarily dictated by two features: (i) properties of the TMD including its hydrophobicity and helical propensity, and (ii) properties of the C-terminal soluble domain that must be translocated across the bilayer during insertion (Kalbfleisch et al., 2007; Costello et al., 2017; Fry et al., 2021). TAs with highly hydrophobic TMDs are preferentially targeted to the ER membrane for insertion via the guided entry of tail-anchored protein (GET) pathway (Schuldiner et al., 2005; Schuldiner et al., 2008). Its central targeting factor in human cells, TRC40 (Stefanovic and Hegde, 2007; Favaloro et al., 2008), binds TMDs using an ordered methionine-rich substrate binding groove and delivers its substrate TAs to the WRB/CAML insertase for membrane integration (Mariappan et al., 2011). TAs with lower hydrophobicity TMDs, however, do not efficiently bind TRC40 and thus, cannot access the GET pathway (Guna et al., 2018). The largest classes of such low hydrophobicity TAs are those targeted for the ER, where they are inserted by the ER membrane protein complex (EMC) (Jonikas et al., 2009; Christianson et al., 2011; Guna et al., 2018), and those targeted to the outer mitochondrial membrane, where they are inserted by MTCH1 and 2 (Guna et al., 2022b). Because of their biophysical similarity, there is thought to be some constitutive levels of mistargeting between these compartments, necessitating dedicated quality control machinery at the ER and mitochondria to extract mislocalized TAs (Chen et al., 2014; Okreglak and Walter, 2014; McKenna et al., 2020).

Since functional constraints limit the potential diversity of the TMD alone, a second sequence element, the short polar C-terminal domain, is known to contribute to TA sorting (Isenmann et al., 1998; Kuroda et al., 1998; Borgese et al., 2007). Though biophysically diverse, mitochondrial TAs are enriched for positive charges in their C-terminal tails, while the C-termini of ER TAs are more likely to be net neutral or negatively charged. Manipulation of C-terminal charge is known to be sufficient to shift the localization of TAs between the ER and mitochondria (Horie et al., 2002; Rao et al., 2016; Costello et al., 2017). However, the biochemical basis for how changes in charge can alter TA sorting is fundamentally not clear. Considering recent advances in our mechanistic understanding of TA insertion into the ER (Pleiner et al., 2020; Bai et al., 2020; O’Donnell et al., 2020; Miller-Vedam et al., 2020), we sought to re-examine the molecular basis for sorting specificity between mitochondrial and ER TAs at this cellular compartment.

## RESULTS

### Selectivity at the ER membrane

Previous studies of the canonical co-translational insertion pathway suggest that sorting fidelity is the combined result of contributions from cytosolic targeting steps and selectivity at the membrane (Trueman et al., 2012; Akopian et al., 2013). In the case of TA proteins, the source of this specificity at either step has remained elusive. While specificity during cytosolic targeting must undoubtedly contribute to TA localization, we found that even when loaded onto the identical chaperone *in vitro*, some mitochondrial TAs cannot be efficiently inserted into the ER membrane (Figure 1A). This observed selectivity appeared to correlate with C-terminal charge, because when positively charged amino acids were introduced within the C-terminus of the canonical ER TA squalene synthase (SQS), its insertion efficiency was dramatically diminished. Based on these observations, we concluded that there must be a source of substrate discrimination directly at the ER membrane, with selectivity occurring at the insertion step.

**Figure 1.**
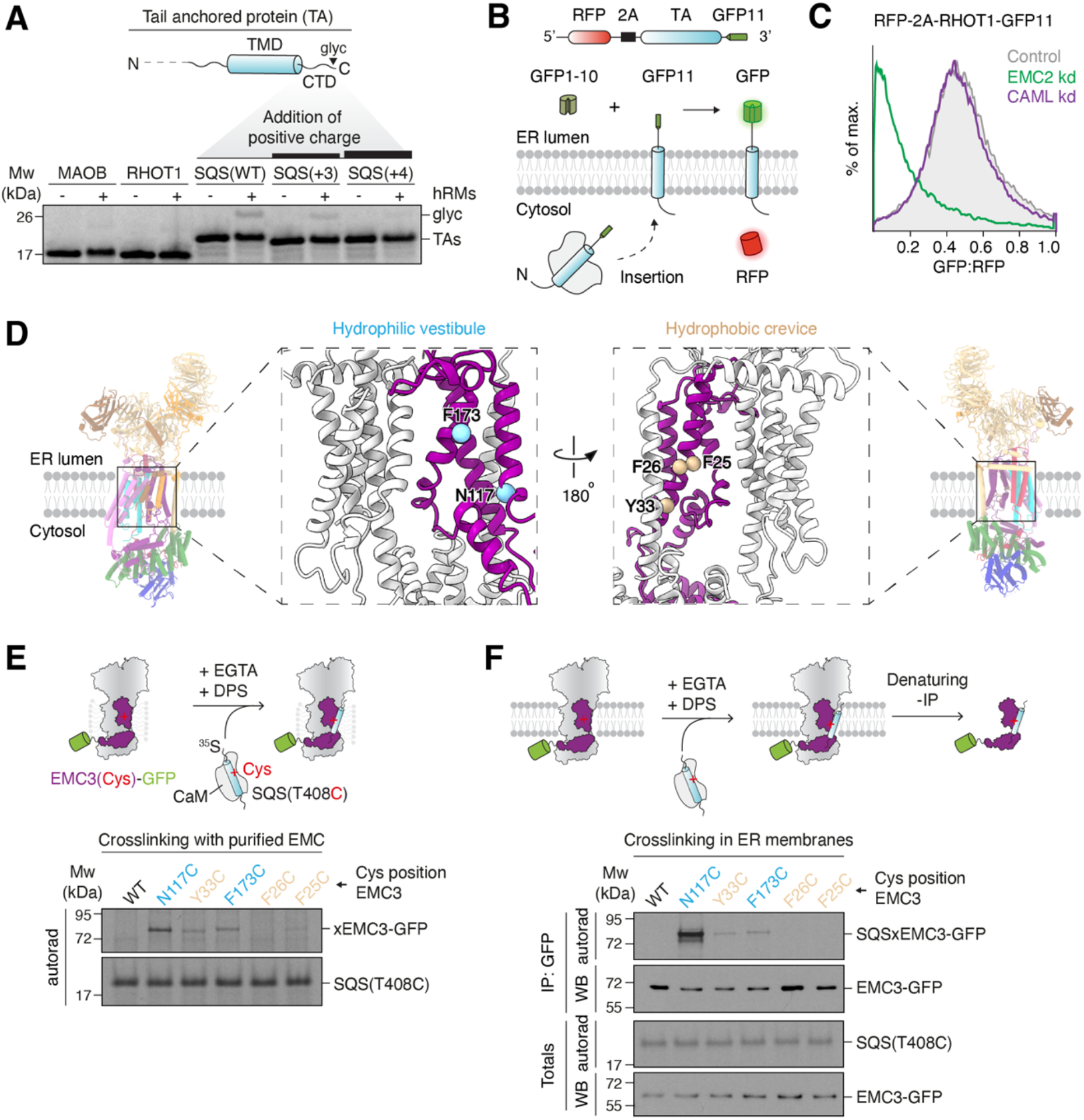
Selectivity at the ER membrane limits misinsertion of mitochondrial TAs by the EMC, which uses a hydrophilic vestibule for insertion. (**A**) (Top) Topology of a TA. (Bottom) ^35^S-methionine-labeled TA reporters with the indicated TMDs and C-terminal domains (CTDs), were expressed in the PURE system and purified as complexes with the cytosolic chaperone calmodulin. Glycosylation (glyc) of its C-terminus upon incubation with human ER microsomes (hRMs) indicates successful insertion. Samples were analyzed by SDS-PAGE followed by autoradiography. (**B**) Schematic of the split GFP reporter system used to selectively monitor TA insertion into the ER. TAs fused to GFP11 are expressed in K562 cells constitutively expressing GFP1-10 in the ER lumen, along with a translation normalization marker (RFP). Successful integration into the ER results in GFP complementation and fluorescence. (**C**) ER insertion of RHOT1-GFP11 as described in (B) was assessed in cells transduced with either a non-targeting (control), EMC2, or CAML knockdown (kd) single guide RNA (sgRNA). GFP fluorescence relative to the normalization marker RFP was determined by flow cytometry and displayed as a histogram. (**D**) Views of the two intramembrane surfaces of the EMC. Residues in EMC3 (purple) lining either the hydrophilic vestibule or hydrophobic crevice were mutated to cysteines for disulfide crosslinking and are highlighted in blue or tan, respectively. EMC4, 7 and 10 are omitted in the inset for clarity. (**E**) Purified wildtype (WT) or EMC3 cysteine (Cys) mutant EMC was incubated with CaM-SQS containing a cysteine at position T408 (CaM-SQS[T408C]). After substrate release from CaM with EGTA, cysteines in close proximity were crosslinked with the zero-length disulfide crosslinker DPS. Quenched reactions were analyzed by SDS-PAGE and autoradiography. (**F**) Human ER-derived microsomes (hRMs) prepared from EMC3 WT or Cys mutant cell lines were mixed with CaM-SQS(T408C) for crosslinking as described in (E). Substrate crosslinks were enriched by denaturing purification of EMC3-GFP. Samples were analyzed by SDS-PAGE followed by autoradiography or western blotting.

The EMC is the major insertase for ER TAs with lower hydrophobicity TMDs, which are similar to those of mitochondrial TAs. Consistent with this biophysical similarity, we and others have demonstrated that the EMC is responsible for misinsertion of mitochondrial TAs into the ER (Figures 1B-C, S1A-B; Guna et al., 2022b; McKenna et al., 2022). Using an established split GFP system to specifically query TA integration into the ER (Inglis et al., 2020), we found that multiple mitochondrial TAs were misinserted in an EMC, but not WRB/CAML, dependent manner (Figures 1C and S1A-B). We therefore reasoned that one source of discrimination against TAs with positively charged C-termini at the ER, either mitochondrial or the SQS mutants, must originate from properties of the EMC.

### Substrate TMDs physically associate with the EMC’s hydrophilic vestibule

With the goal of determining the biochemical basis of EMC’s substrate specificity, we sought to map the path of a TMD from the cytosol into the bilayer through the EMC. Structures of the yeast and mammalian EMC identified two intramembrane surfaces that could potentially catalyze TMD insertion: a hydrophilic vestibule that positions several conserved positively charged residues within the cytosolic leaflet of the bilayer, and a hydrophobic crevice that contains a large lipid-filled wedge within the membrane (Pleiner et al., 2020; Bai et al., 2020; O’Donnell et al., 2020; Miller-Vedam et al., 2020). Site-specific crosslinking experiments previously identified EMC3 as the major substrate interaction partner within the purified EMC (Pleiner et al., 2020), consistent with EMC3’s homology with other members of the Oxa1 superfamily of insertases (Anghel et al., 2017). However, the path of a substrate TMD has never been directly determined, and potential contributions to insertion from both intramembrane surfaces of the EMC have been proposed.

To map direct physical association of substrates with the EMC, we exploited several independent zero-length crosslinking approaches to chart substrate interaction at single-residue resolution. First, we introduced the site-specific crosslinker BpA into the TMD of a canonical TA substrate, and identified UV-dependent crosslinks to both EMC3 and EMC4 by immunoprecipitation (Figure S1C). Unlike EMC3, which is present on both sides of the complex, the cytosolic and intramembrane surfaces of EMC4 partially enclose only the hydrophilic vestibule, suggesting substrates must at least transiently localize with this side of the EMC. Second, we exploited the fact that endogenous EMC3 does not contain any naturally occurring cysteine residues to perform disulfide crosslinking between a TA and the EMC. Because disulfide-bond formation can only occur between residues within 3-5 Å of each other, productive crosslinking necessarily indicates a direct physical association. Zero-length disulfide formation between single cysteines introduced at defined positions in EMC3, and a unique cysteine within a substrate TMD, identified a strong preference for substrate binding to the hydrophilic vestibule of detergent-solubilized EMC (Figures 1D-E and S1D). A similar preference was observed when comparing matched positions on either side of EMC3 at the base of the membrane. This preferential crosslinking was independent of cysteine position within the substrate TMD (Figure S1E) and was also observed upon incorporation of orientation-independent photo-crosslinkers in EMC3 (Figure S1F).

Finally, and most definitively, we developed a strategy to capture the transient interaction between a substrate TMD and the EMC by disulfide crosslinking in native, insertion competent, ER membranes (Figures 1F and S1G-H). Using this approach, we again observed a marked preference for interaction of TAs with the hydrophilic vestibule of EMC3 compared to the hydrophobic crevice. In native membranes and with purified EMC, substrates preferentially crosslinked to a cytosol-facing position on EMC3 at the entrance to the lipid bilayer, suggesting a potential increase in dwell time at this location.

To further exclude that the opposite hydrophobic crevice is involved in TA insertion, we introduced multiple mutations to polar and hydrophobic residues in this region and found that they are all dispensable for TA biogenesis in human cells (Figure S2). These data, in combination with sequence conservation, homology to Oxa1 superfamily insertases, and mutational analysis, definitively identify the hydrophilic vestibule as the insertase competent module of the EMC.

### An improved model of the EMC defines intramembrane surfaces required for insertion

Having identified the hydrophilic vestibule as the major site of substrate binding to the EMC, we sought to better define its architecture and thereby identify potential sources of substrate specificity. The insertase core of the EMC (composed of EMC3 and 6) is partially enclosed by the dynamic subunits EMC4, 7 and 10. However, whether EMC7 and 10 contain TMDs, how these may be positioned, as well as the specific contributions of all three auxiliary subunits was incompletely defined.

To characterize the biophysical properties of the hydrophilic vestibule we obtained an improved cryoelectron microscopy (cryo-EM) reconstruction of the human EMC that allowed us to unambiguously assign and position the three TMDs of EMC4 and the single TMDs of EMC7 and 10 (Figures 2A, S3 and S4A-C; Table 1). In support of this model, we biochemically confirmed that human EMC7 and 10 both contain single C-terminal TMDs that span the lipid bilayer (Figure S4D). Examination of the roles of these subunits suggested that, consistent with previous studies, EMC4 and 7, but not 10 are required for TA biogenesis (Figure S4E; Louie et al., 2012; Volkmar et al., 2019; Lakshminarayan et al., 2020). These auxiliary subunits do not play an architectural role in complex stability, as their depletion did not affect assembly of the core EMC subunits (EMC2,3,5,6,8) (Figure S4F). However, we additionally found that complete loss of EMC4 impaired the assembly of EMC7 and 10 into the EMC. Because EMC4’s C-terminal β-strand completes the membrane-proximal β-propeller of EMC1, it is possible that loss of EMC4 disrupts the lumenal binding sites of EMC7 and 10. We concluded that the hydrophilic vestibule formed by the TMDs and cytosolic loops of EMC3 and 6 is partially enclosed by the five dynamic TMDs of EMC4, 7 and 10.

**Figure 2.**
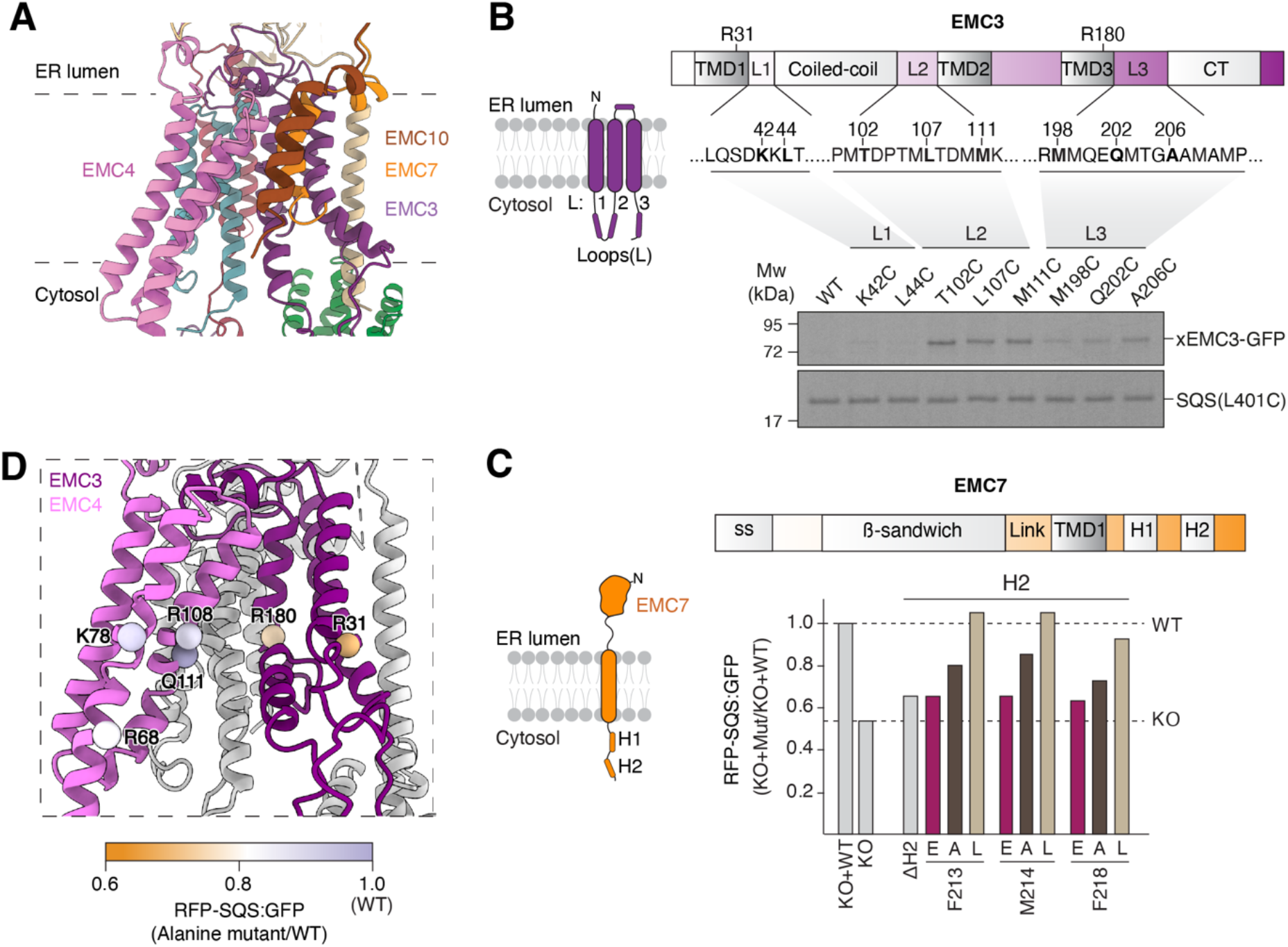
Characterization of the cytosolic and intramembrane residues required for insertion by the EMC. (**A**) Displayed is an improved model of the human EMC determined using cryoelectron microscopy (cryo-EM), of the insertase core, composed of EMC3/6, enclosed by the three TMDs of EMC4, and the single TMDs of EMC7 and 10. (**B**) (Top) Schematic of the topology and domain organization of EMC3, highlighting three flexible cytosolic loops (L1-3) located beneath the hydrophilic vestibule of the EMC. (Bottom) Purified wildtype (WT) or EMC3 Cys mutant EMC were incubated with purified CaM-SQS(L401C) complexes for disulfide crosslinking and analysis as in Figure 1E. (**C**) (Top) Schematic of the topology and domain organization of EMC7. ss = signal sequence; Link = linker; H1 = helix 1; H2 = helix 2. (Bottom) HEK293 EMC7 knockout (KO) cells were transduced with lentivirus to express WT EMC7, or the indicated mutants of EMC7 helix 2 (H2). The effects of each mutant on biogenesis of SQS was determined using the ratiometric fluorescent reporter assay, normalized to WT and plotted as a bar chart. (**D**) View of the hydrophilic vestibule with EMC7 and 10 omitted for clarity. Residues indicated with spheres are colored according to the effects of individual alanine mutations at these positions in EMC3 and 4 on expression of SQS in HEK293T cells. The effect of each mutant was determined by flow cytometry using the ratiometric fluorescent reporter assay in (C), normalized to wildtype, and is displayed according to the indicated legend.

**Table 1.** Cryo-EM data collection, refinement, and validation statistics

|  | Consensus<br>(EMDB-xxxx) | Nine-subunit<br>(EMDB-xxxx)<br>(PDB xxxx) | Eight-subunit<br>(EMDB-xxxx) |
| --- | --- | --- | --- |
| **Data collection and processing** |  |  |  |
| Microscope | FEI Titan Krios |  |  |
| Voltage (kV) | 300 |  |  |
| Camera | Gatan K3 |  |  |
| Magnification | 105,000 |  |  |
| Defocus range (μm) | -1.0 to -3.0 |  |  |
| Pixel size (Å/pix) | 0.416 |  |  |
| Electron exposure (e-/Å<sup>2</sup>) | 60 |  |  |
| Number of frames per movie | 40 |  |  |
| Dose Rate (e-/pix/s) | 16.0 |  |  |
| Automation software | SerialEM |  |  |
| Number of micrographs | 11,822 |  |  |
| Initial particle images (no.) | 1,271,124 |  |  |
| Final particle images (no.) | 193,900 | 156,706 | 37,194 |
| Local resolution range | 3.0 – 7.0 | 3.0 – 7.0 | 3.5 – 8.0 |
| Map resolution range (Å, FSC=0.143) | 3.5 | 3.6 | 3.9 |
| **Refinement** |  |  |  |
| Software (phenix.real_space_refine) | PHENIX 1.20.1-4487 |  |  |
| Initial model used (PDB code) |  | 6WW7 + Alpha Fold |  |
| Correlation coefficient (CC<sub>mask</sub>) |  | 0.82 |  |
| Map sharpening *B* factor (Å<sup>2</sup>) | -112 | -103 | -76 |
| Model composition |  |  |  |
| Non-hydrogen atoms |  | 17,837 |  |
| Protein residues |  | 2259 |  |
| Ligands |  | NAG:5 PCW |  |
| *B* factors (Å<sup>2</sup>) |  |  |  |
| Protein |  | 105.42 |  |
| Ligand |  | 90.63 |  |
| R.M.S deviations |  |  |  |
| Bond lengths (Å) (# > 4σ) |  | 0.010 |  |
| Bond angles (°) (# > 4σ) |  | 0.961 |  |
| **Validation** |  |  |  |
| MolProbity score |  | 1.99 |  |
| Clashscore |  | 15.51 |  |
| Poor rotamers (%) |  | 0.48 |  |
| Cβ deviations (%) |  | 0.05 |  |
| CaBLAM outliers (%) |  |  |  |
| Ramachandran plot |  |  |  |
| Favored (%) |  | 95.69 |  |
| Allowed (%) |  | 4.27 |  |
| Disallowed (%) |  | 0.04 |  |

### Capture of substrate TAs in the cytosol by the EMC

Based on this improved model of the EMC, we determined that the cytosolic loops of EMC3 and 7 are positioned immediately below the hydrophilic vestibule, making them prime candidates for cytosolic capture of substrates. We had previously shown that the flexible loops of EMC3 contain conserved methionine residues, commonly found in the TMD binding domains of cytosolic chaperones, that were important for EMC function (Pleiner et al., 2020). We therefore hypothesized that the loops of EMC3 and 7 could be involved in physically interacting with substrate TMDs in the cytosol. We set out to test key facets of this working model, with the goal of understanding whether the molecular details of substrate capture could contribute to discrimination between ER and mitochondrial TAs.

Consistent with earlier data, we found that methionine residues within the cytosolic loop of EMC3 were essential for TA biogenesis in cells (Figures 2B and S5A-C). Similarly, we found that the flexible C-terminus of EMC7 was required for EMC function (Figures 2C and S6A-D). Deletion of twelve residues to disrupt a predicted amphipathic α-helix, but not deletion of a matched upstream α-helix, strongly impaired SQS biogenesis, nearly phenocopying EMC7 knockout. We further demonstrated that the hydrophobicity of conserved residues within both this amphipathic helix of EMC7 and the methionine-rich loops of EMC3 is important, because their mutation to leucine, but not alanine or glutamate supported wild type levels of EMC function in cells (Figure 2C,D). However, for these loops to be directly involved in TA capture, they must be capable of physically interacting with substrate TMDs. Indeed, using zero-length disulfide crosslinking, we found that the cytosolic loops of EMC3 and 7 specifically interact with substrates in a TMD dependent manner (Figures 2B and S6E-F).

We concluded that the primary role of these flexible loops is to position hydrophobic residues within the cytosol, which physically capture substrate TMDs for subsequent insertion into the membrane. Because the biophysical properties of the TMDs of mitochondrial and ER targeted TAs are in many cases indistinguishable, there was no clear mechanism by which capture by EMC3 and 7 could contribute to substrate discrimination based on C-terminal charge. We therefore turned to consideration of the intramembrane surfaces of the hydrophilic vestibule.

### Substrates must passage through a positively charged hydrophilic vestibule for insertion

The improved atomic model of the EMC enabled detailed structure-function analysis of the biophysical requirements of the hydrophilic vestibule for TA insertion. The defining characteristic of the hydrophilic vestibule is a network of conserved polar and positively charged residues within the cytosolic leaflet of the lipid bilayer. Previous analysis suggests that charged and polar residues required for EMC function are positioned within the TMDs of the core insertase subunits EMC3 and 6 (Pleiner et al., 2020). Mutations to the positively charged residues in EMC3 strongly impaired insertion in cells, whereas mutations to EMC6 had only mild effects.

A more complete understanding of the localization of EMC4, 7 and 10 allowed us to systematically introduce mutations to all of the polar residues that face the EMC3/6 insertase core (Figure 2D). However, we found that mutations to polar, charged, and methionine residues within EMC4’s TMDs had little to no effect on TA biogenesis (Figures 2D and S7A-C). Only mutation of residues that likely affect TMD packing (N140) or lipid headgroup interaction (K67) showed significant phenotypes. If EMC4 does not directly contribute to function, it may instead be playing a role in regulating access to the hydrophilic vestibule, as deletion of its cytosolic EMC2-binding site strongly impaired SQS biogenesis (Figure S7D-E). Of all the polar intramembrane residues tested within the hydrophilic vestibule, the highly conserved R31 and R180 of EMC3 are the most crucial for TA insertion, and their combined mutation displayed an additive effect on substrate biogenesis (Figures 2D and S7F-G).

### Positively charged soluble domains impede insertion by the EMC

Both these mutational data and our crosslinking results together suggest that substrates must passage into the membrane directly along a positively charged surface of EMC3. Mislocalization of a mitochondrial TA into the ER requires both insertion of its TMD and translocation of its associated positively charged C-terminal domain. Thus, we reasoned that the positively charged hydrophilic vestibule is ideally positioned to discriminate mitochondrial and ER TAs through charge repulsion (Figure 3A).

**Figure 3.**
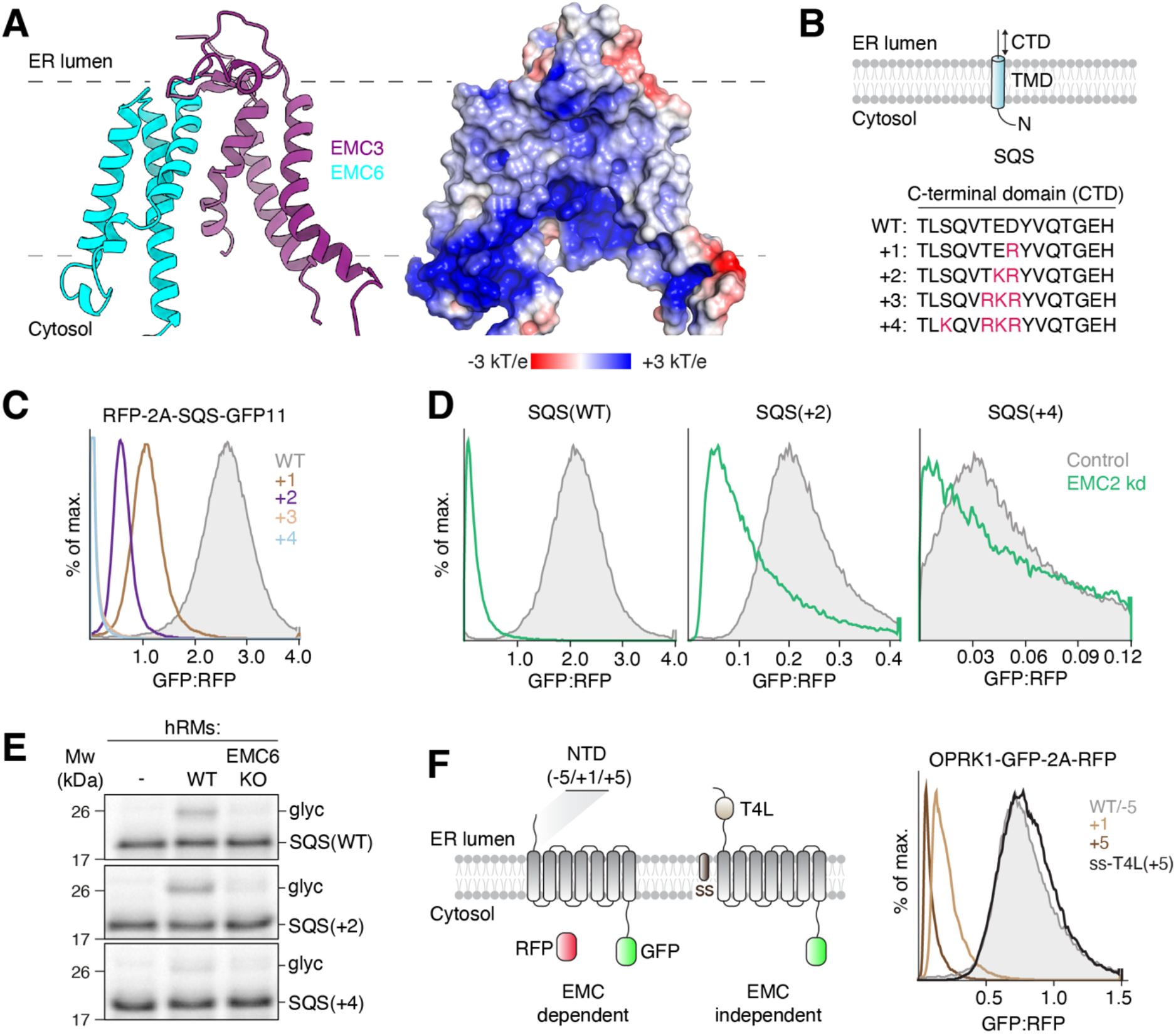
Positively charged soluble domains impair EMC insertion. (**A**) (Left) Model of the TMDs of EMC3 and 6 forming the insertase core. (Right) Surface representation of the electrostatic potential of the insertase core ranging from −3 to +3 kT/e. EMC4, 7, and 10 were omitted for clarity. (**B**) Schematic of the SQS C-terminal domain (CTD) charge series. The C-terminus of SQS was mutated to introduce positively charged residues at the indicated positions. (**C**) Integration of the indicated SQS mutants into the ER as determined using the split GFP reporter system described in Figure 1B. (**D**) As in (C), but for cells expressing either a non-targeting (control) or EMC2 knock down (kd) sgRNA. (**E**) The indicated ^35^S-methionine labeled SQS charge mutants were expressed in rabbit reticulocyte lysate and incubated with human ER-derived microsomes (hRMs) prepared from HEK293 wildtype (WT) or EMC6 knockout (KO) cells. ER insertion is monitored by glycosylation (glyc) of an acceptor motif fused to the C-terminus of the TA substrates. (**F**) WT (−5) or the indicated N-terminal domain (NTD) charge mutants of the GPCR opioid receptor kappa 1 (OPRK1) GFP-fusions were expressed along with an RFP normalization marker in RPE1 cells. Cells were analyzed by flow cytometry and the GFP:RFP ratio is displayed as a histogram. Bypassing insertion by the EMC by fusion to a cleavable signal sequence (ss) enhances ER integration of the OPRK1(+5) charge mutant.

To test the fundamental premise of this hypothesis, we first characterized the impact of charge on insertion by the EMC. In order to directly query the role of C-terminal charge, without confounding effects from comparing different substrates or TMDs, we generated a series of mutants of the canonical ER TA, SQS, containing increasing amounts of positive charge within its soluble C-terminal domain (Figure 3B). Using the split GFP reporter system, we found that while all SQS mutants inserted into the ER in an EMC-dependent manner, insertion efficiency was inversely correlated with positive charge (Figures 3C-D and S8A). Even addition of a single positive charge to the C-terminus of SQS resulted in a dramatic decrease in integration into the ER. Validating that this effect is specifically occurring at the insertion step and cannot be explained by other effects in cells (e.g. substrate stability), we observed a similar trend between charge and insertion into ER microsomes *in vitro* (Figures 3E and S8B).

In addition to its role in TA insertion, the EMC co-translationally inserts the first Nexo TMD of many GPCRs (Chitwood et al., 2018). Like the C-termini of ER TAs, the N-termini of GPCR are typically short, unstructured, and de-enriched for positive charges (Figure S8C) (Wallin and von Heijne, 1995). Using the EMC-dependent GPCR OPRK1, we found that introduction of positive charge is again inversely correlated with insertion propensity by the EMC (Figures 3F and S8D). We therefore propose that inefficient translocation of positively charged extracellular domains is an inherent property of the EMC shared by both its co- and post-translational insertase function.

### A selectivity filter in the EMC contributes to TA protein sorting fidelity

The EMC’s strong bias against translocation of positively charged domains provides a biochemical explanation for discrimination of mitochondrial TAs at the ER. To determine if this selectivity is due at least in part to charge repulsion between the hydrophilic vestibule of the EMC and the soluble C-terminal domain of a substrate TA, we tested whether manipulation of the electrostatic potential of the EMC could alter substrate selectivity.

Due to the prominent location of R31 and R180 of EMC3 at the cytosolic entrance to the hydrophilic vestibule, these residues are ideally positioned to form a charge barrier that selectively prevents translocation across the lipid bilayer. If true, mutations that alter the electrostatic potential of these residues could alleviate repulsion between the EMC and positively charged soluble domains, allowing increased misinsertion of mitochondrial TAs. Mutation of both EMC3 R31 and R180 to alanine or glutamate did not affect EMC assembly, and as expected markedly impaired insertion of SQS in cells using our ratiometric fluorescent reporter system (Figures 4A and S9A-C). However, SQS variants containing increasingly positively charged C-termini showed increased insertion by the glutamate, but not the alanine mutant EMC. A similar trend was observed for insertion of SQS variants *in vitro* into wild type, alanine, or glutamate mutant ER microsomes, validating that charge specifically affects insertion propensity (Figures 4B and S9D-E). Similarly, these EMC3 mutations differentially affected the insertion of the co-translational substrate OPRK1 and its positively charged N-terminal domain mutants in cells (Figure S9F).

**Figure 4.**
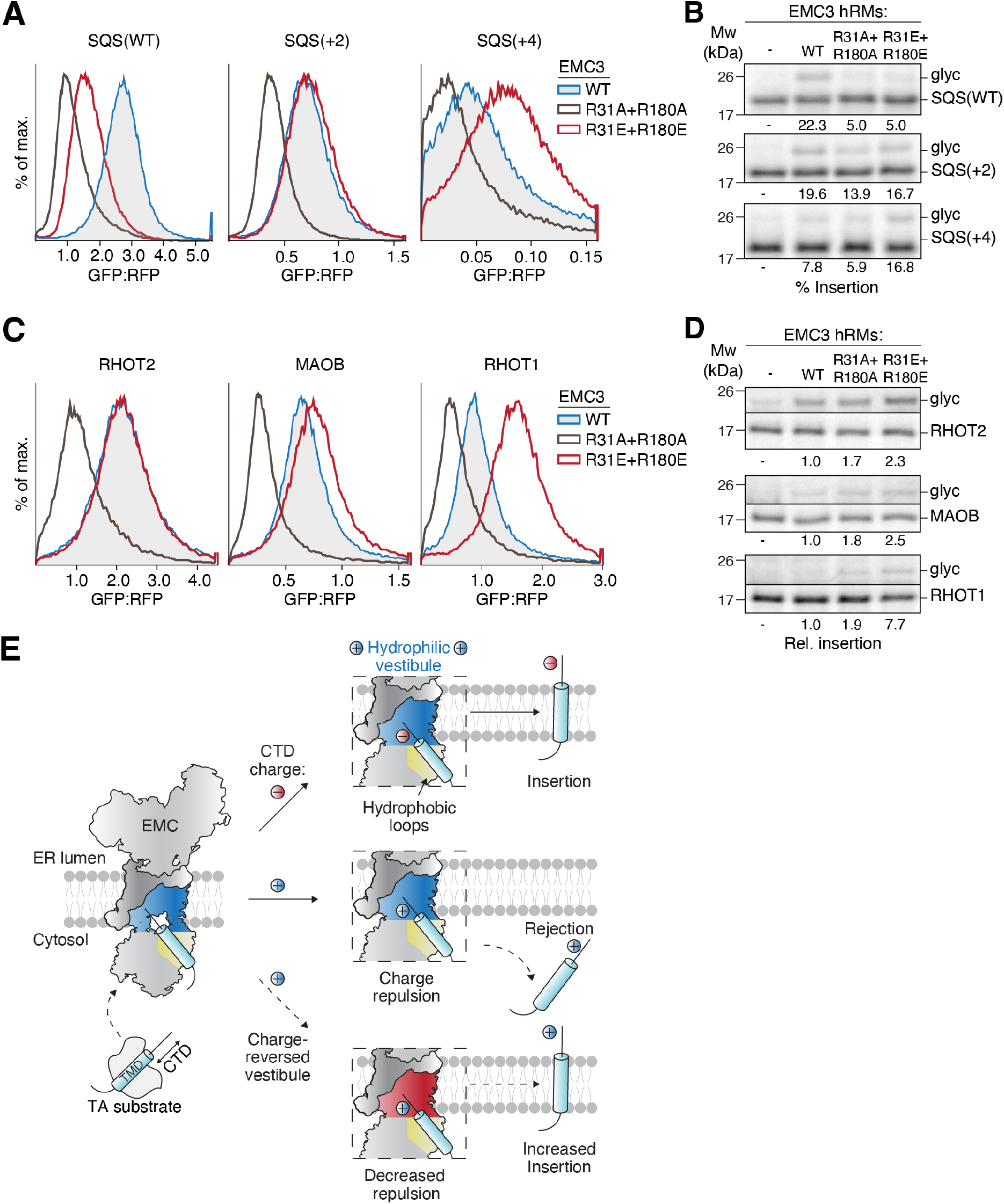
A selectivity filter in the EMC limits mitochondrial TA misinsertion at the ER. (**A**) ER insertion of the indicated SQS charge mutants was measured in cells expressing either wild type (WT), R31A+R180A, or R31E+R180E EMC using the split GFP reporter system described in Figure 1B. (**B**) The indicated SQS mutants were prepared as in Figure 3E, incubated with hRMs from EMC3 WT, R31A+R180A or R31E+R180E expressing cell lines. Successful ER insertion is monitored with a glycosylation (glyc) acceptor motif fused to the C-terminus of each substrate. The % glycosylated is indicated below the gel. (**C**) As in (A), but with the indicated mitochondrial TAs. Note the strong increase in ER mis-localization of RHOT1 in EMC3 R31E+R180E expressing cells. (**D**) As in (B) but expressing the TMD and C-terminus of the indicated mitochondrial TAs in non-nucleased rabbit reticulocyte lysate. (**E**) Model for how the EMC distinguishes TA clients by C-terminal charge. TA TMDs are initially captured by flexible hydrophobic loops in the cytosol, allowing their C-terminus to probe the net positively charged hydrophilic vestibule. In the absence of positive charge, the C-terminus is translocated rapidly, enabling TMD insertion. Insertion of TAs with positively charged C-termini is slowed by charge repulsion, which facilitates TA dissociation (rejection). Charge repulsion can be alleviated by introducing negative charge into the hydrophilic vestibule, resulting in increased misinsertion.

Finally, because these SQS variants serve only as a proxy for the effects of charge on insertion, we tested whether manipulation of the EMC selectivity filter could also affect mislocalization of bona fide mitochondrial TAs into the ER. Indeed, we found that multiple mitochondrial TAs, most notably RHOT1, showed increased ER insertion upon expression of the glutamate, but not the alanine mutant EMC3 in cells and *in vitro* (Figures 4C-D and S9G). Fis1, MAOA and MAOB similarly showed increased ER insertion.

## DISCUSSION

These results suggest that charge repulsion at the EMC provides a selectivity filter to control the subcellular localization of TAs (Figure 4E), enforcing their accurate sorting between the ER and mitochondrial outer membrane. The enrichment of positive charge in the C-termini of mitochondrial (and likely peroxisomal) TAs, serves as a flag for discrimination at the ER by the EMC. Unlike its TMD, which must mediate function and targeting, the C-terminal domain of most TAs is functionally dispensable and may have evolved primarily to facilitate sorting specificity. The combined evolution of mitochondrial TA’s positively charged C-termini and the positively charged hydrophilic vestibule of the EMC thereby limits misinsertion of TAs at the ER membrane.

The molecular basis for TA discrimination was revealed by a systematic analysis of substrate insertion *in vitro* and in cells that defines the path through the hydrophilic vestibule of the EMC into the membrane. After delivery to the ER by a cytosolic chaperone, the first step in substrate insertion is handover and capture by the EMC. We found that substrate TMDs physically interact with the conserved, hydrophobic loops of EMC3 and EMC7 located immediately beneath the vestibule in the cytosol. Mutational analysis suggests that only the hydrophobicity of these loops, but not their specific amino acid sequence, is important for TA insertion. Indeed, comparison of EMC3 with its bacterial and archaeal homologs suggests that methionine-rich cytosolic loops are a conserved feature of Oxa1 superfamily insertases (Borowska et al., 2015), but the specific positioning of these hydrophobic residues is not strictly critical. We propose that these hydrophobic loops represent the first transient, flexible interaction site for substrate TMDs by the EMC.

We observed that substrates crosslink more efficiently to both these loops and the cytosol-exposed residues of the hydrophilic vestibule than to residues within the lipid bilayer. This difference was especially pronounced in native insertion-competent membranes, more likely to represent on-pathway intermediates that are not artefacts of detergent solubilization. These data would be consistent with a longer dwell time of substrates in this cytosolic intermediate followed by faster partitioning into the lipid bilayer. Similarly, a recent kinetic analysis of the bacterial insertase YidC suggests it undergoes rapid substrate capture via its cytosolic loops and substantially slower translocation of the polar domain and membrane insertion (Laskowski et al., 2021). A plausible explanation for this observation might be that translocation of a polar domain across the hydrophobic lipid bilayer has a high energetic barrier and thus is a rate-limiting step to insertion.

This would be consistent with molecular dynamics simulations that suggest that TMD partitioning into the membrane is an energetically favorable process and membrane protein insertases are primarily required to decrease the energetic barrier for translocation of a soluble domain across the bilayer (Nicolaus et al., 2021; White and Wimley, 1999). Therefore, interaction of a substrate TMD with EMC’s cytosolic hydrophobic loops could prevent aggregation, while its C-terminus probes the hydrophilic vestibule. For correctly targeted TAs, the EMC’s hydrophilic vestibule serves as a funnel that catalyzes translocation of their C-termini into the ER lumen by providing a hydrophobicity gradient between the aqueous cytosol and the core of the bilayer. Positioning of similar hydrophilic grooves or vestibules within a locally thinned membrane is a common feature of evolutionary distinct protein translocases (Kumazaki et al., 2014; Voorhees et al., 2014; McDowell et al., 2020; Wu et al., 2020), and represents a striking example of convergent evolution. In the case of the EMC, the dynamic TMDs of EMC4, 7 and 10 provide a protected environment, devoid of any potential off-pathway interaction partners, for the nascent protein to sample the bilayer.

However, for mistargeted mitochondrial or peroxisomal TAs, the positive net charge of the hydrophilic vestibule would impose a kinetic barrier to translocation of their positively charged C-terminal domains. In these TAs, positive charges are frequently found clustered near their TMD, suggesting that charge density and positioning, rather than simple net charge, determines the extent of charge repulsion at the EMC. Repulsion likely delays translocation and thus increases the chance of TA dissociation from the hydrophobic loops. Using purified components, we previously showed that the cytosolic domain of the EMC does not contain an ordered high-affinity TMD binding site (Pleiner et al., 2021), as can be found in Get3 or SRP (Guna and Hegde, 2018). A composite transient TMD capture surface formed by flexible hydrophobic loops might allow for faster dissociation of TA clients and thus enable quicker accept/reject decisions. Rejected TAs in the cytosol could then be either recaptured for targeting to the correct organelle, or triaged for degradation by quality control machinery. In this way, the EMC directly contributes to the accurate sorting of the ~600 TAs that must be expressed and localized to distinct organelles in human cells.

The two positively charged residues in EMC3, which provide the charge barrier for entrance to the hydrophilic vestibule, are universally conserved in all Oxa1 superfamily insertases. As a result, its homologs, including WRB and YidC, have also been suggested to inefficiently translocate positively charged soluble domains (Rao et al., 2016; Soman et al., 2014). Indeed, the effect of charge on insertion efficiency appears to be an inherent quality of the EMC and affects both its post- and co-translational substrates. Similar to EMC’s TA substrates, GPCRs that lack an N-terminal signal sequence and are therefore potential EMC clients, typically contain neutral or negatively charged N-terminal extracellular domains (Figure S8C; Wallin and von Heijne, 1995). Using the same strategy for discrimination of mitochondrial TAs, the EMC could also enforce the positive inside rule (von Heijne, 1986) for co-translational substrates by limiting translocation of positively charged domains into the ER lumen.

In summary, we have characterized the molecular logic for how the EMC contributes to selective membrane protein localization in human cells. Its function is analogous to the active role Sec61 plays in substrate selection and rejection at the ER (Trueman et al., 2012). Whether MTCH1 and 2 also confer similar contributions to substrate selectivity at the mitochondrial outer membrane is an important question for future research. However, specificity at the membrane is only one layer of the multi-faceted approach used to regulate protein sorting. Cells employ a sieved strategy in which the overall fidelity of protein localization is the combined result of selectivity at each biogenesis step from chaperone binding in the cytosol to insertion at the membrane (Rao et al., 2016). Particularly in metazoans, where membrane protein mislocalization can lead to disease (Juszkiewicz and Hegde, 2018), these steps are tightly coupled to quality control machinery that ensures immediate recognition and degradation of failed intermediates. By limiting misinsertion of TAs and potentially preventing topological errors in multipass membrane proteins, the EMC serves as a guardian for protein biogenesis at the ER.

## Acknowledgement

We thank Songye Chen and Oliver Clarke for technical assistance, all members of the Voorhees lab for thoughtful discussion, and Alina Guna for critical reading of the manuscript. We thank Pamela Bjorkman for access to her lab’s cell sorter, as well as the Caltech Flow Cytometry facility, and the Caltech Cryo-EM facility. Cryo-electron microscopy was performed in the Beckman Institute Center for TEM at Caltech, and data was processed using the Caltech High Performance Cluster, supported by a grant from the Gordon and Betty Moore Foundation. This work was supported by: the Heritage Medical Research Institute (RMV), the NIH’s National Institute Of General Medical Sciences DP2GM137412 (RMV), the Deutsche Forschungsgemeinschaft (TP), and the Tianqiao and Chrissy Chen Institute (TP, MH).

## MATERIALS & METHODS

### Plasmids and antibodies

Constructs for *in vitro* translations in rabbit reticulocyte lysate were based on the pSP64 vector (Promega, USA). Constructs for *in vitro* translation in the *E. coli* PURExpress system were generated from the T7 PURExpress plasmid (New England Biolabs, USA). pSpCas9(BB)-2A-Puro (PX459) and lentiCRISPR v2 were gifts from Feng Zhang (Addgene plasmids #48139 and #52961). pLG1-puro non-targeting sgRNA 3, used for cloning CRISPRi sgRNAs, was a gift from Jacob Corn (Addgene plasmid #109003). The 2nd generation lenti-viral packaging plasmid psPAX2 was a gift from Didier Trono (Addgene plasmid #12260) and the envelope plasmid Δ8.9 was a kind gift from Carlos Lois. The pHAGE2 lenti-viral transfer plasmid was a gift of Magnus A. Hoffmann and Pamela Bjorkman. For expression in K562 cells, a lenti-viral backbone containing a UCOE-EF-1α promoter and a 3’ WPRE element was used (Addgene #135448), which was a kind gift of Martin Kampmann and Jonathan Weissman. The expression plasmid for the SENP^EuB^ protease (Addgene ID #149333) was a gift of Dirk Görlich. Plasmids for amber suppression in mammalian cells were kind gifts of Simon Elsässer. Note that the mCherry variant of RFP was used throughout this study, but the simpler nomenclature of RFP is used in the text and figures. Similarly, EGFP is used throughout this study, but referred to as GFP.

The following antibodies were used in this study: EMC2 (25443-1-AP, Proteintech, USA); EMC3 (67205-1-Ig, Proteintech, USA); EMC4 (27708-1-AP, Proteintech, USA); EMC5 (A305-833, Bethyl Laboratories, USA); EMC7 (27550-1-AP, Proteintech, USA); EMC10 (ab180148, Abcam, UK); CAML (Synaptic Systems, Germany); Anti-HA-HRP (Millipore-Sigma, USA); Anti-FLAG-HRP (Millipore-Sigma, USA). The rabbit polyclonal antibodies against BAG6 and GFP were gifts from Ramanujan Hegde. Secondary antibodies used for Western blotting were: Goat anti-mouse- and anti-rabbit-HRP (#172-1011 and #170-6515, Bio-Rad, USA).

The following sgRNAs were cloned into PX459 or lentiCRISPR v2 and used to generate knock out cell lines: EMC3 (AAGAAAGTGATGATAACGAT); EMC4 (TCATACACACCATCATAGTA); EMC6 (GCCGCCTCGCTGATGAACGG); EMC7 (TTCTCCGTCTACCAGCACTC); EMC10 (AGTGCCAACTTCCGGAAGCG). The following sgRNAs were cloned into pLG1 for CRISPRi knockdowns: non-targeting control (GGCTCGGTCCCGCGTCGTCG); EMC2 (GCCATCTTCCCAGAACCTAG); CAML (ATGTTGGCCGCCGCTGCGA).

### Expression and purification of biotinylated anti-GFP and anti-ALFA nanobody

Protease-cleavable biotinylated anti-GFP and anti-ALFA tag nanobodies (Götzke et al., 2019; Kirchhofer et al., 2010) that were used for EMC purifications throughout this study were expressed in *E. coli* and purified using Ni^2+^-chelate affinity chromatography using protocols described in detail before (Pleiner et al., 2015; Pleiner et al., 2020). The expression of His_14_-Avi-SUMO^Eu1^-anti GFP nanobody from plasmid pTP396 (Addgene #149336) was carried out with the following modification. Instead of biotinylating the nanobody *in vitro* with purified biotin ligase BirA, pTP396 was expressed in the *E. coli* strain AVB101 (Avidity, USA), which contains an IPTG-inducible plasmid for BirA co-expression. 50 μM biotin was added to the main culture 1 h before induction of nanobody and BirA expression.

The sequence of the ALFA^ST^ nanobody was derived from the original study describing its generation (Götzke et al., 2019) and cloned into pTP396. Expression was carried out in *E. coli* Rosetta-gami 2 cells (Millipore-Sigma, USA) in a 1 L scale for 6 h at 18°C after induction of protein expression with 0.2 mM IPTG. The resulting His_14_-Avi-SUMO^Eu1^-anti ALFA nanobody fusion protein was purified from cell lysate using Ni^2+^-chelate affinity chromatography for *in vitro* biotinylation with purified biotin ligase BirA as described before (Pleiner et al., 2020).

Immobilized biotinylated nanobodies were cleaved off of streptavidin magnetic beads using an engineered SUMO protease (SENP^EuB^) that recognizes the SUMO^Eu1^ module (Vera Rodriguez et al., 2019). His_14_-Tev-tagged SENP^EuB^ protease (Addgene ID #149333) was expressed in *E. coli* NEB express I^q^ as described before (Pleiner et al., 2020). For sequential immunoprecipitations, a commercial system with orthogonal cleavage sites based on the SUMOStar tag and SUMOStar protease (LifeSensors, USA) (Liu et al., 2008) was used.

### Conjugation of ALFA nanobody to HRP for Western blotting

To use the ALFA nanobody in Western blotting, it was coupled to HRP-maleimide via a single engineered C-terminal cysteine residue as described for other nanobodies before (Pleiner et al., 2018).

### Mammalian *in vitro* translation

*In vitro* translation reactions in rabbit reticulocyte lysate (RRL) were carried out with *in vitro* transcribed mRNA as described before (Sharma et al., 2010). PCR products generated from pSP64-derived plasmids or gene fragments (synthesized by Integrated DNA Technologies or Twist Biosciences, both USA) served as templates for run-off transcription and contained a 5’ SP6 promoter followed by an open-reading frame and a 3’ stop codon. A 10 μl transcription reaction contained 7.6 μl T1 mix (Sharma et al., 2010), 0.2 μl SP6 polymerase (New England Biolabs, USA), 0.2 μl RNAsin (Promega, USA), 100 ng PCR product, and was carried out for 1.5 h at 37°C. Transcriptions were added directly to RRL. Unless indicated otherwise, RRL was treated with S7 micrococcal nuclease (Roche, Germany) in the presence of CaCl2 to remove endogenous hemoglobin mRNA. Nascent proteins are labeled during translation reactions of 15-30 min at 32°C in RRL by incorporation of radioactive ^35^S-methionine (Perkin Elmer, USA). Nascent TA proteins were released from the ribosome with 1 mM puromycin and then incubated with 5% (v/v) of either canine pancreatic rough microsomes (cRMs) (Walter and Blobel, 1983) or human ER-derived microsomes (hRMs), prepared from engineered cell lines as described below, for another 20 min at 32°C. Samples were analyzed by SDS-PAGE and autoradiography to detect the translated ^35^S-labeled TA protein.

Successful post-translational insertion into microsomes was monitored by glycosylation of a canonical NXS/T acceptor motif. This was appended either as part of a charged C-terminal opsin tag (MNGTEGPNFYVPFS<u>**N**KTVD</u>) or where no additional C-terminal domain charge was desired, an NGT motif was placed 22 amino acids downstream of the TMD after a neutral glycine-serine linker and followed by an additional C-terminal GS dipeptide.

### Protease protection assay

To assess the membrane spanning topology of EMC7 and EMC10, they were tagged with an N-terminal 1xHA and a C-terminal 3xFLAG tag and translated in RRL in the presence of cRMs as described above. Protease-accessible regions of both proteins were digested by incubation with 0.5 mg/ml Proteinase K for 1 h at 4°C in the presence or absence of 0.05% (v/v) Triton-X-100 to solubilize cRM membranes. Proteinase K was inactivated by addition of 5 mM PMSF and quick transfer into boiling SDS buffer (100 mM Tris/HCl pH 8.4; 1% [w/v] SDS). Denatured digestion reactions were diluted tenfold with IP buffer (50 mM HEPES/KOH pH 7.5; 300 mM NaCl; 0.5 % [v/v] Triton-X-100) and incubated with anti-HA or anti-FLAG M2 resin (Millipore-Sigma, USA) for 1 h at 4°C for immunoprecipitation of protected fragments.

### Preparation of human ER-derived microsomes (hRMs)

To prepare hRMs from Expi293 suspension cell lines, cells were harvested and then washed twice in 50 ml 1x PBS. Cells were then resuspended in 4x pellet volume of sucrose buffer (10 mM HEPES/KOH pH 7.5.; 2 mM MgAc; 250 mM Sucrose, 1x Protease inhibitor cocktail [Roche, Germany]) and lysed with ~50 strokes in a tight-fit dounce homogenizer. Complete cell lysis was verified by trypan blue staining. The lysate was then diluted twofold and spun for 30 min at 3214 g in a table-top centrifuge at 4°C to remove nuclei and cell debris. This spin was repeated and the resulting supernatant was then centrifuged for 1 h at 75,000 g at 4°C (TLA-100.3 rotor or Type60 Ti rotor; Beckman Coulter, USA). The supernatant was aspirated and the membrane pellet gently resuspended in microsome buffer (10 mM HEPES/KOH pH 7.5.; 1 mM MgAc; 250 mM Sucrose, 0.5 mM DTT). Membranes prepared for disulfide crosslinking were resuspended in microsome buffer without DTT. The absorbance at 280 nm of the resuspended membranes was measured by boiling an aliquot in SDS buffer (100 mM Tris/HCl pH 8.4; 1% [w/v] SDS). The hRM preparation was then adjusted to an absorbance of 75 at 280 nm using microsome buffer. To remove endogenous mRNAs, the adjusted hRM preps were further treated with S7 micrococcal nuclease (Roche, Germany) at a concentration of 0.075 U/μl in the presence of 0.33 mM CaCl_2_ for 6 min in a 25°C water bath, then quickly removed to ice and quenched by Ca^2+^-chelation with 0.66 mM EGTA. Nucleased hRMs were snap-frozen in liquid nitrogen in single-use aliquots and stored until further use at −80°C.

### *In vitro* translation of TA proteins in the PURExpress system

Plasmids containing a 5’ T7 promoter, followed by an open-reading frame, stop codon and 3’ T7 terminator were used as templates for the coupled *in vitro* transcription/translation PURExpress system (New England Biolabs, USA). The various SQS constructs used for cysteine crosslinking comprised an N-terminal 3xFLAG tag, the human Sec61β cytosolic linker (residues 2-59) with the natural cysteine at position 39 mutated to serine, as well as the five N-terminal flanking residues, TMD and complete C-terminus of human FDT1/SQS (residues 378-end). Cysteine residues were introduced at the indicated positions using site-directed mutagenesis. TA translations were supplemented with radioactive ^35^S-methionine and 10 μM purified Calmodulin (CaM) (Shao et al., 2017).

For use in photocrosslinking reactions, TA substrates were generated that contained the unnatural amino acid and photocrosslinker 4-Benzoylphenylalanine (BpA) (Bachem, Switzerland), which was incorporated into the TMD by amber stop codon suppression in the PURExpress system lacking all release factors (ΔRF123; New England Biolabs, USA). The release factors RF2 and RF3, but not RF1 (which recognizes the UAG [amber] stop codon) were added back to the reaction. BpA was added at 100 μM and incorporated at UAG codons using purified BpA aminoacyl-tRNA synthetase and suppressor tRNA, prepared as described before (Shao et al., 2017).

All PURE translation reactions were carried out for 2 h at 32°C and then ribosome-associated nascent chains were released by addition of 1 mM puromycin (Thermo Fisher Scientific, USA) and further incubation for 10 min at 32°C. To remove aggregated protein, the translation reactions were layered over a 20% (w/v) sucrose cushion prepared in physiological salt buffer (PSB) (50 mM HEPES/KOH pH 7.5; 130 mM KAc, 2 mM MgAc) that further contained 100 nM CaCl_2_. After a 1 h spin at 55,000 rpm (TLS-55 rotor; Beckman-Coulter, USA) at 4°C, soluble TA-CaM complexes were retrieved from the supernatant.

### Photocrosslinking

Purified EMC complexes in detergent micelles for photocrosslinking were obtained via anti-GFP nanobody IP from stable human suspension cell lines that ectopically expresses GFP-EMC2. They were mixed with ^35^S-Methionine labeled BpA-containing TA-CaM complexes generated in the PURExpress system as described above. TAs were released from CaM shortly before UV radiation by addition of 1 mM EGTA to chelate calcium. Except for the -UV control sample, all reactions were irradiated at a distance of ~7-10 cm with a UVP B-100 series lamp (Analytik Jena, Germany) for 15 min on ice before quenching with SDS-PAGE sample buffer. Samples were adjusted to 1% (w/v) SDS and boiled. Denatured reactions were diluted tenfold with IP buffer (50 mM HEPES/KOH pH 7.5; 300 mM NaCl; 0.5 % [v/v] Triton-X-100) and incubated with Protein A sepharose beads (Thermo Fisher Scientific, USA) and EMC3 or EMC4 antibodies for immunoprecipitation. Samples were analyzed by SDS-PAGE and autoradiography.

Site specific incorporation of the photocrosslinking amino acid 3’-azibutyl-N-carbamoyl-lysine (AbK) into EMC3 in mammalian cells was performed by amber suppression using the *Methanosarcina mazei* pyrrolysyl-tRNA synthetase (PylRS)/tRNA^Pyl^CUA (PylT) pair (Ai et al., 2011). Constructs for amber suppression in mammalian cells were created as follows using previously reported plasmids as template (Elsässer et al., 2016). The first plasmid encodes 4 copies of PylT(U25C), as well as WT PylRS, which was further modified by mutating Y306A and Y384F to accommodate the bulky AbK (Yanagisawa et al., 2008; O’Donnell et al., 2020). The coding region of EMC3 was inserted with a C-terminal GFP-tag into a second plasmid which also encoded 4 additional copies of PylT(U25C). Selected amino acid positions in EMC3 were mutated to amber stop codons, for incorporation of AbK at these sites. To generate AbK-containing EMC, Expi293 cells (Thermo Fisher Scientific, USA) were transiently co-transfected with 4xPylT/PylRS(Y306A, Y384F) and 4xPylT/EMC3(Amber[TAG])-GFP plasmids at a ratio of 4:1 using PEI “MAX” (Polysciences, USA). The cells were grown in the presence of 0.5 mM AbK (Iris Biotech, Germany) and harvested 72 h after transfection. EMC complexes with successfully suppressed Amber stop codons, contained full length AbK-modified EMC3 and could thus be purified via the C-terminal GFP-tag as described below. The purified EMC complexes were mixed with ^35^S-Methionine labeled SQS(WT)-CaM complexes generated in the PURExpress system and irradiated with UV as described above. Samples were analyzed by SDS-PAGE and autoradiography.

### Disulfide crosslinking

EMC complexes containing wild type or cysteine mutant EMC3 or EMC7 variants were purified from stable human suspension cell lines as described below and mixed with wild type or cysteine mutant SQS-CaM complexes generated in the PURExpress system as described above.

The zero-length disulfide crosslinker 4,4’-Dipyridyldisulfide (DPS) (Millipore-Sigma, USA) was added to a final concentration of 250 μM to initiate the crosslinking of cysteines in close proximity after SQS release from CaM with 1 mM EGTA. The reaction was incubated for 2 h on ice and analyzed by SDS-PAGE and autoradiography.

For disulfide crosslinking in membranes, hRMs were prepared from stable human suspension cell lines expressing wild type or cysteine mutant EMC3 variants as described above. hRMs were mixed with PURE translated SQS-CaM complexes in PSB and 500 μM DPS. After substrate release with 500 μM EGTA, reactions were incubated for 2 h on ice before quenching with 5 mM L-Cysteine (Millipore-Sigma, USA). The reactions were then adjusted to 1% (w/v) SDS and incubated at room temperature for 10 min to denature the EMC complex. The denatured reactions were diluted tenfold with IP buffer (50 mM HEPES/KOH pH 7.5; 300 mM NaCl; 0.5 % [v/v] Triton-X-100) and the EMC3-GFP subunit was specifically enriched via the GFP tag. After elution by boiling in sample buffer containing 0.5 M urea, the samples were analyzed by SDS-PAGE and autoradiography.

### Cell culture and cell line generation

Adherent HEK293 cell lines were cultured in Dulbecco’s Modified Eagle’s Medium (DMEM) supplemented with 10% fetal calf serum (FCS) and 2 mM L-Glutamine. For Flp-In T-Rex 293 cell lines containing integrated doxycycline-inducible reporters, tetracycline-free FCS was used and culture medium additionally supplemented with 15 μg/ml blasticidin S and 100 μg/ml hygromycin B. RPE1 cells were cultured in DMEM/F-12 (1:1) supplemented with 10% FCS and 2 mM L-Glutamine.

Flp-In 293 T-Rex cells were purchased from Thermo Fisher Scientific (USA). Stable Flp-In 293 T-Rex cell lines designated as GFP-2A-RFP-SQS/VAMP2 express the RFP-tagged transmembrane domain and flanking regions of human squalene synthase (SQS/FDFT1) or vesicle-associated membrane protein 2 (VAMP2). The generation of these cell lines was described previously (Guna et al., 2018; Pleiner et al., 2020). In these cell lines, GFP is expressed as a soluble cytosolic protein from the same mRNA as RFP-SQS/VAMP2 using a viral 2A sequence that induces peptide-bond skipping by the ribosome (de Felipe et al., 2006). Their RFP and GFP fluorescence intensity can be measured by flow cytometry to derive a RFP:GFP ratio. Changes in this ratio after perturbation, e.g. expression of a mutant EMC subunit, reflect differences in the post-translational stability of the TA reporter.

The stable, doxycycline-inducible GFP-EMC2 Flp-In 293 T-Rex cell line and its adaptation to suspension growth in FreeStyle 293 Expression Medium (Thermo Fisher Scientific, USA) was described before (Pleiner et al., 2020). Clonal knockouts of EMC4, 7 and 10 in this background were obtained by transfecting the adherent parental cell line with PX459 encoding the respective sgRNA using TransIT-293 transfection reagent (Mirus, USA). 48 h post transfection, 1 μg/ml puromycin was added for three consecutive days. Medium was subsequently exchanged to allow for two days of recovery before single cell clones were seeded into 96-well plates by limiting dilution. Knockout efficiency of the selected clones was verified by Western blotting and the resulting adherent knockout cell lines were either used directly for flow cytometry experiments or adapted to suspension growth for EMC purifications.

Expi293 cells (Thermo Fisher Scientific, USA) were maintained at a concentration of 0.5-2.0 million cells per ml in Expi293 Expression Medium (Thermo Fisher Scientific, USA). An EMC3 knockdown suspension cell line was generated by transient transfection of Expi293 cells with an EMC3 sgRNA cloned into lentiCRISPR v2 using PEI “MAX” (Polysciences, USA). Transfected cells were treated with 10 μg/ml puromycin for four consecutive days. Then the medium was exchanged to allow for 10 days of recovery. The polyclonal cell population demonstrated a sufficient level of consistent downregulation of endogenous EMC3 and was thus used directly to re-introduce wild type EMC3 or various mutants tagged with a C-terminal TagBFP or GFP via lenti-viral transduction as described below. Transduced cell lines were sorted using fluorescence of the fused TagBFP or GFP to obtain a homogenous population of cells with near full replacement of endogenous EMC3 with a tagged mutant copy of interest. Wild type EMC7 or various cysteine mutants with an N-terminal ALFA tag were introduced via lenti-viral transduction into the EMC3-GFP cell line.

A K562 CRISPRi cell line, stably expressing dCas9-BFP-KRAB Tet-ON (Jost et al., 2017), was transduced with lentivirus as described below to constitutively express β-strands 1-10 of superfolder GFP (residues 2-214) (Cabantous et al., 2005) in the ER lumen via fusion to an N-terminal signal sequence and a C-terminal KDEL sequence as described previously (Guna et al., 2022b).

### CRISPRi knockdowns

K562 dCas9-BFP-KRAB Tet-ON, ER GFP1-10 cells were transduced via spinfection as described below with lentivirus containing a pLG1-puro backbone and a sgRNA targeting a gene of interest. Sequences of sgRNAs were derived from the hCRISPRi-v2 compact library (Horlbeck et al., 2016). 48 h after spinfection, 1 μg/ml puromycin was added for three consecutive days to select cells with a successfully integrated sgRNA expression cassette. After two days of recovery, cells were transduced with GFP11-tagged TA reporters expressed from a lentiviral backbone under control of a UCOE-EF1α promoter. Cells were analyzed 48 h after reporter spinfection by flow cytometry (8 days after sgRNA transduction).

### Lenti-viral transduction

Lentivirus was generated by co-transfection of HEK293T cells with a desired transfer plasmid and two packaging plasmids (psPAX2 and Δ8.9) using the TransIT-293 transfection reagent (Mirus, USA). 48 h post transfection, culture supernatant was harvested, aliquoted and flash frozen in liquid nitrogen.

For transduction of Expi293 or suspension-adapted Flp-In 293 T-Rex cells, 20 million cells were mixed with 2.5 ml freshly harvested lenti-viral supernatant (i.e. the complete supernatant from one 6-well of lenti-producing HEK293T cells 48 h after transfection) in 20 ml medium in a 125 ml vented Erlenmeyer flask (Celltreat, USA). Then the flask was transferred to a shaking incubator and transduced cells were grown for around 16 hours. Cells were then pelleted, resuspended in 50 ml of fresh medium and grown for 2-3 days before sorting of successfully transduced cells on a SH800S cell sorter (Sony Biotechnology, USA).

K562 cells were transduced by spinfection. Briefly, 250,000 cells were mixed with 50-200 μl of lentiviral supernatant and RPMI medium in the presence of 8 μg/ml polybrene in a total volume of 1 ml in a 24-well plate. 24-well plates were spun at 1,000 xg for 1.5 h at 30°C. Cells were then resuspended and transferred to a 6-well plate. Lenti-viral reporter constructs used in K562 cells for flow cytometry analysis all contained an upstream UCOE-EF1α promoter, followed by RFP, a P2A site and the full length human coding regions for all Mito TAs fused to GFP11 via a five residue Gly-Ser linker. SQS mutants were expressed in the same cassette, but contained the cytosolic linker (residues 2-70) of human Sec61β at the N-terminus followed by the TMD, N-terminal flanking region and complete C-terminus of of human FDFT1/SQS (residues 378-417 [end]). Charge mutations were introduced as shown in Figure 3B. EMC3 WT or its arginine mutants were expressed in K562 cells from a lentiviral transfer plasmids with an upstream EF1α promoter and fused to a C-terminal TagBFP-3xFLAG tag.

For lenti-viral transduction of adherent HEK293 or RPE1 cells, 50-200 μl lentiviral supernatant and 8 μg/ml polybrene (Millipore-Sigma, USA) were usually added directly to ~70% confluent cells in 2.5 ml culture medium in a 6-well. Lenti-viral reporter constructs of SQS and VAMP2 for use in HEK293 cells (Figures 2C, S4E, S6B-C, S7D) contained an upstream CMV promoter, followed by GFP, a 2A site and RFP, which was directly fused to the TMD and flanking regions of human FDFT1/SQS or VAMP2 as described before (Guna et al., 2018; Pleiner et al., 2020). OPRK1 reporter constructs used in RPE1 cells, expressed full length human OPRK1 (WT/-5), OPRK1(E45K,D46R,E50K)(+1 variant) or OPRK1(E35K,D37R,E45K,D46R,E50K) (+5 variant) as N-terminal fusions to EGFP, followed by a 2A site and RFP from a CMV promoter.

### Flow cytometry analysis of reporter cell lines

All adherent cells were trypsinized, washed, and resuspended in 1xPBS for flow cytometry analysis. K562 cells were analyzed directly. Analysis was either on an Attune NxT Flow Cytometer (Thermo Fisher Scientific, USA) or a MACSQuant VYB (Miltenyi Biotec, Germany). Flow cytometry data was analyzed using FlowJo v10.8 Software (BD Life Sciences, USA). Unstained cells transiently transfected with either GFP, RFP (or BFP if needed) were analyzed separately along every run as single-color controls for multicolor compensation using the FlowJo software package.

For experiments in K562 cells, lenti-viral fluorescent reporters were introduced via spinfection as described above usually 48 h before analysis. To probe the effect on EMC2 or CAML knockdown on reporter insertion, cells were additionally transduced with sgRNA expressing lenti-viral vectors as described under ‘CRISPRi knockdowns’. To analyze the effect of EMC3 mutations on TA reporters, K562 cells were first spinfected with lentivirus expressing EMC3(WT/mut)-BFP. After 48 h, mitochondrial TA or SQS charge mutant reporter lentivirus was spinfected. Cells were analyzed by flow cytometry after another 48 h. Adherent HEK293 or RPE1 cells were analyzed 48 h after transduction as described above.

### Purification of engineered EMCs from stable suspension cell lines

Stable human suspension cell lines expressing tagged wild type or mutant copies of EMC subunits were generated and grown as described above. EMC complexes were purified using anti-GFP or anti-ALFA nanobody essentially as described before (Pleiner et al., 2015; Pleiner et al., 2020). Cells were harvested by centrifugation for 10 min at 3,000 g and washed in 1xPBS. Cell pellets were resuspended with 6.8 ml solubilization buffer (50 mM HEPES/KOH pH 7.5; 200 mM NaCl; 2 mM MgAc; 1% [w/v] LMNG, 1 mM DTT, 1x complete EDTA-free protease inhibitor cocktail [Roche, Germany]) per 1 g of cell pellet and incubated for 30 min at 4°C. Lysates were cleared by centrifugation for 30 min at 4°C at 18,000 rpm (SS-34 rotor; Beckman-Coulter, USA).

In parallel, Pierce magnetic Streptavidin beads (Thermo Fisher Scientific, USA) were equilibrated in wash buffer (solubilization buffer with 0.0025% [w/v] LMNG) and then incubated with biotinylated anti-GFP or anti-ALFA tag nanobody, purified as described above. After nanobody immobilization, free biotin binding sites were blocked by incubation with wash buffer containing 10 μM biotin-PEG-COOH (Iris Biotech, Germany) for 10 min on ice. Blocked, nanobody-decorated beads were then added to cell lysate for binding to detergent-solubilized ALFA- or GFP-tagged EMC complexes for 1 h at 4°C with head-over-tail mixing. Magnetic beads were then collected and washed three times with wash buffer, before resuspension of the beads in wash buffer containing 250 nM SENP^EuB^ protease in a volume amounting to one half of the original bead suspension volume. Protease elution was allowed to proceed for 20 min on ice.

EMC complexes containing fully replaced cysteine mutant EMC7 variants, were purified via a 2-step procedure using first the C-terminal GFP tag on EMC3 and then the N-terminal ALFA tag on EMC7. The GFP nanobody eluate, obtained by SENP^EuB^ cleavage, was diluted twentyfold with wash buffer and incubated with beads containing immobilized ALFA nanobody. The ALFA nanobody was tagged with an orthogonal SUMOStar protease cleavage site and bound EMC was then eluted along with the ALFA nanobody in wash buffer containing 500 nM SUMOStar protease. The resulting eluate was aliquoted and flash frozen in liquid nitrogen. The concentrations of purified EMC complexes for disulfide crosslinking were normalized by measuring GFP fluorescence on a BioTek Synergy HTX plate reader (Agilent, USA). Normalization was verified by SDS-PAGE and Sypro Ruby staining (Thermo Fisher Scientific, USA). If necessary, normalizations were adjusted based on the quantification of Sypro Ruby stained EMC subunit bands in Fiji.

### Purification of EMC for structure determination

A suspension-adapted GFP-EMC2 Flp-In 293 T-Rex cell line (Pleiner et al., 2020) was used to purify the EMC for structural analysis. Additionally, EMC7 carrying a C-terminal ALFA tag was introduced into this cell line via lenti-viral transduction as described above. The lenti-viral transfer plasmid encoded EMC7-ALFA fused via a viral 2A sequence to BFP (EMC7-ALFA-2A-TagBFP). BFP fluorescence was used to sort a homogenous stable suspension cell line that ectopically expresses both GFP-EMC2 and EMC7-ALFA. EMC was purified as described above, but with the following minor modifications. Cells were solubilized with solubilization buffer containing 1% glyco-diosgenin (GDN) (Anatrace, USA). The wash buffer contained 0.05% [w/v] GDN. Finally, the EMC eluate was concentrated using an Amicon Ultra 0.5 ml 100K MWCO concentrator (Millipore-Sigma, USA) and further purified via size-exclusion chromatography using a Superose 6 Increase 3.2/300 column (Cytiva, USA) equilibrated in wash buffer (50 mM HEPES/KOH pH 7.5; 200 mM NaCl; 2 mM MgAc; 0.05% [w/v] GDN and 1 mM DTT). Fractions corresponding to the EMC were pooled and concentrated as above to 0.5 mg/ml. To reduce the conformational flexibility of EMC7 at the insertase side, we added stoichiometric amounts of purified ALFA nanobody (Götzke et al., 2019), which binds the C-terminal ALFA tag on EMC7.

### Grid preparation and data collection

CryoEM grids were prepared by applying 3 μl of purified EMC at 0.5 mg/mL to glow discharged (60 seconds using a Pelco easiGlow, Emeritech K100X at a plasma current of 20 mA), Holey carbon grids (Quantifoil R1.2/1.3). The sample was blotted for 4-6 sec with filter paper at 8°C, 100% humidity at a −4-blot force prior to plunging into liquid ethane for vitrification using the FEI Vitrobot Mark v4 x2 (Thermo Fisher Scientific, USA). The data set was acquired on a Titan Krios electron microscope (Thermo Fisher Scientific, USA) operated at 300 keV equipped with a K3 direct electron detector and an energy filter (Gatan, USA) with a 20-eV slit width. A total of 11,822 micrographs were collected using 3-by-3 pattern beam image shift, acquiring movies for three non-overlapping areas per hole, using an automated acquisition pipeline in SerialEM (Mastronarde, 2005). Movies were recorded with 40 frames at a magnification of 105,000x in super resolution mode at a calibrated magnification of 0.416 Å/pixel using a dose of 60 e-/Å2 at a dose rate of 16.0 e-/pixel/s and a defocus range of −1.0 to −3.0μm.

### Image processing

The data processing workflow is summarized in Figure S3 and was performed using cryoSPARC v.3.3–v.4.0 (Punjani et al., 2017). In short, 11,822 micrographs were motion corrected, dose weighted and down sampled (two-fold to 0.832 Å/pixel) using the Patch Motion followed by patch-based CTF estimation using Patch CTF. 10,206 movies were selected and manually curated using cut-offs for CTF fit (5.0 Å) and total motion (50 pix) for further processing. The particle picking was done using the automated Blob Picker function with particle diameter of 150 to 400 Å. After two rounds of 2D classification, 1,271,124 particles were used for two rounds of heterogeneous ab initio reconstruction (4 volumes), using Maximum/Initial resolution of 9 and 7Å respectively and an Initial/Final minibatch size of 400 and 1,200 particles respectively. Once we obtained an initial map with clear features of the EMC, we reclassified the 1.2 million particles using 3D heterogeneous classification using one well-defined class of the EMC and three decoy classes, using a batch size of 5,000 particles per class and initial low-pass filter of 50 Å. Prior to the final round of classification of 212,440 particles were re-extracted in a box size of 400 pix. The final round of classification yielded a population 193,900 particles that were further refined using non-uniform refinement to obtain a reconstruction at 3.5 Å resolution.

To explore the previously observed flexibility between the lumenal, membrane, and cytoplasmic domains, the particles were subjected to two rounds of 3D-variability analysis/clustering, selecting five modes and a filter resolution ranging from 4.0-8.0 Å. After carefully analyzing each reconstruction, a mode corresponding to a missing subunit of the EMC was identified. The subset of particles was then split into 20 clusters using 3D Variability Analysis Display for this mode. Particles belonging to the nine-subunit complex (156,706 particles) that contained high-resolution features were combined and refined using non-uniform refinement. This yielded a map with a resolution of 3.6 Å, in which we detected a stronger EM density for the TMDs of EMC4 and 7.

Particles belonging to the eight-subunit complex (37,194 particles) were combined and similar to the nine-subunit complex, the particles were refined using non-uniform refinement. This yielded a map with a resolution of 3.9 Å. All three maps (consensus, 9- and 8-subunit) were post-processed by applying a sharpening B factor of −112 Å^2^, −103 Å^2^ and −76 Å^2^, respectively. Finally, for the analysis of EMC10’s TMD position a low-pass filter of 5.5 Å was applied to each map using volume tools in cryoSPARC.

All map resolutions were calculated at the final round of refinement using the gold standard FSC=0.143 criterion from the half maps. Statistics details of the EMC EM maps are reported in Table 1.

### Model building and refinement

An initial model for the nine-subunit EMC was generated by docking the EMC structure in a lipid nanodisc (PDB: 6WW7) (Pleiner et al., 2020) into the cryo-EM density using UCSF Chimera (Pettersen et al., 2004) followed by an initial round of refinement using Phenix (Liebschner et al., 2019). Next, for the not well-ordered TMDs of EMC4 and 7 high-confidence subcomplexes EMC3 (residues 5-42 and 101-209), EMC4 (59-155), EMC6 (12-end) and EMC7 (155-178) were generated using AlphaFold2—Multimer ColabFold (AlphaFold2_advanced.ipynb) (Mirdita et al., 2022) and then rigid body fitted into the densities. Finally, all models were combined and further manual refinement was conducted in COOT (Casañal et al., 2019; Emsley et al., 2010). Next, lipids, N-glycans and disulfide bond pairs were added where justified by both the EM density and its chemical environment. Finally, the final model was refined against the 9-subunit map using phenix.real_space_refine. Although we could successfully model a backbone through the contiguous density of the TMDs of EMC4 and 7, we could not unambiguously assign its registry and therefore these TDMs were assigned as poly-Ala/Gly in the final model. Statistics detailed of the EMC model are reported in Table 1. Figures were made using PyMol (Schrödinger LLC), and UCSF ChimeraX.

## SUPPLEMENTARY FIGURES

**Figure S1.**
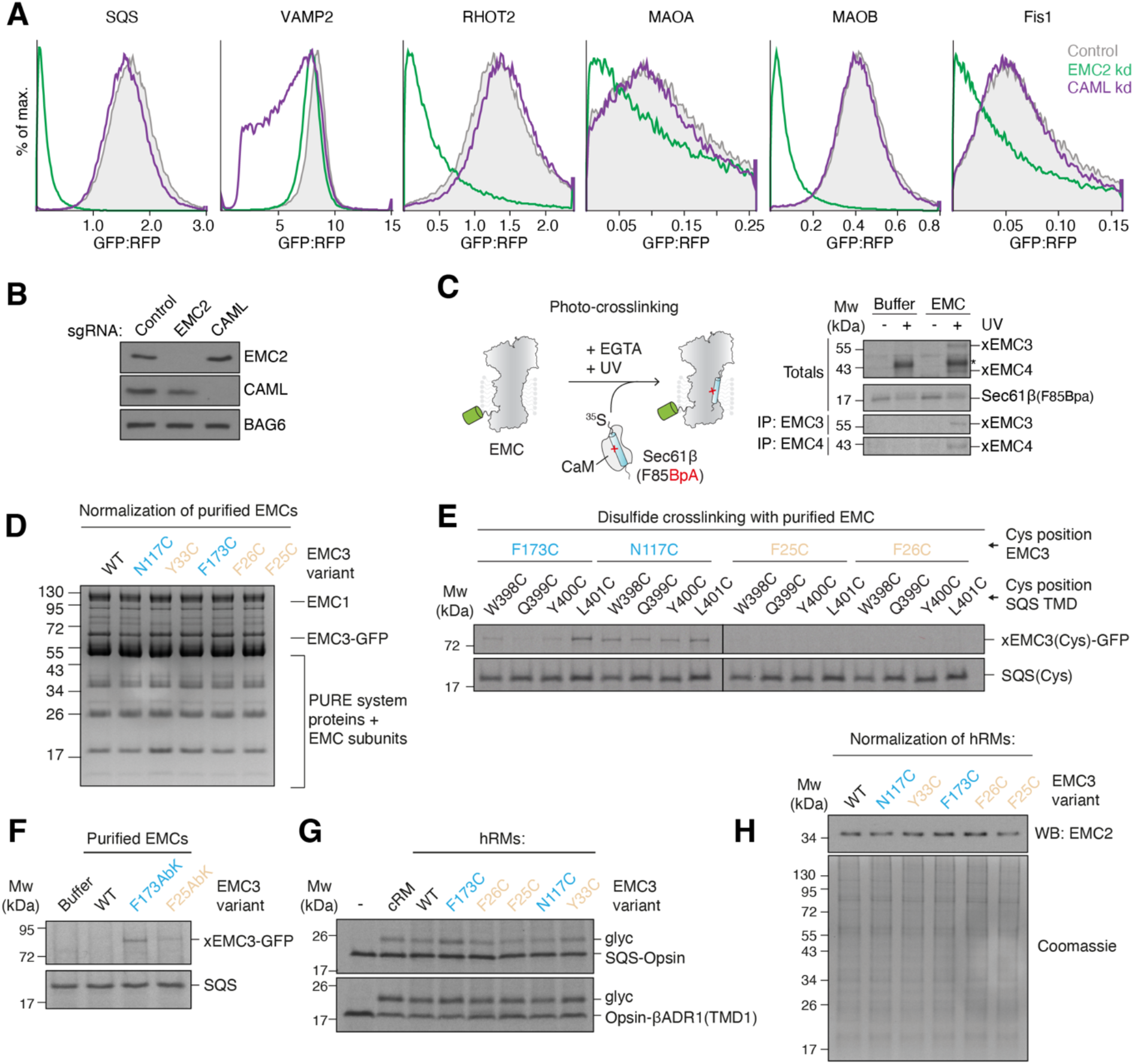
Determining the physical association of substrates with the EMC by site-specific crosslinking. (**A**) Flow cytometry analysis of ER insertion of the indicated ER (SQS, VAMP2) and mitochondrial (RHOT2, MAOA, MAOB, Fis1) TAs using the split GFP system as described in Figure 1B. K562 cells additionally expressed non-targeting (control), EMC2 or CAML knock down (kd) sgRNAs. The effect on ER insertion, relative to an expression control, was determined by flow cytometry and displayed as a histogram. (**B**) Cells from (A) were harvested and samples of total cell lysates were analyzed by SDS-PAGE and western blotting with antibodies against EMC2, CAML and BAG6, a non-targeted control protein. (**C**) Schematic depiction of the site-specific photocrosslinking approach. The ^35^S-methionine labeled TA substrate SEC61β, with a BpA photo–crosslinker incorporated into its TMD, was produced as a complex with calmodulin (CaM) in the PURE *in vitro* translation system. It was then incubated with purified EMC solubilized and purified in the detergent LMNG. Except for the -UV controls, all reactions were irradiated with UV light after substrate release from CaM with EGTA and then analyzed by SDS-PAGE and autoradiography. Crosslinks to EMC3 and EMC4 were identified by immunoprecipitation (IP) with anti-EMC3 and -EMC4 antibodies. The asterisk indicates the crosslinked TA dimer band. (**D**) Coomassie stained SDS-PAGE gel of the disulfide crosslinking experiment with the purified EMC shown in Figure 1E before analysis via autoradiography. The gel shows that equal amounts of EMC were used in the different crosslinking reactions. (**E**) Disulfide crosslinking with purified EMC as in Figure 1E, but with cysteines positioned around a turn of the SQS TMD, showing that the observed crosslinking bias to residues on the hydrophilic vestibule (in blue) is independent of cysteine position. All crosslinking reactions were performed in parallel, and gels were exposed to the same film. (**F**) Purified EMC complexes containing the unnatural amino acid and photocrosslinker Abk incorporated into EMC3 at the indicated positions were mixed with SQS(WT)-CaM complexes prepared in the PURE system and irradiated with UV light after substrate release from CaM with EGTA. Samples were analyzed by SDS-PAGE and autoradiography. (**G**) Insertion activity of human ER-derived microsomes (hRMs) prepared from EMC3 WT or Cys mutant cell lines. Two well-characterized EMC substrates, SQS and TMD1 of the β-adrenergic receptor 1 (βADR1) (Chitwood et al., 2018; Guna et al., 2018), were translated in rabbit reticulocyte lysate in the presence of the indicated hRMs. Successful ER insertion results in the glycosylation (glyc) of the fused opsin tag. Canine pancreatic rough microsomes (cRMs) were used as a control. (**H**) The totals of the disulfide crosslinking reactions in hRMs shown in Figure 1F were analyzed by Western blotting for EMC2 and Coomassie staining of total protein content to show similar amounts of hRMs were used in the experiment.

**Figure S2.**
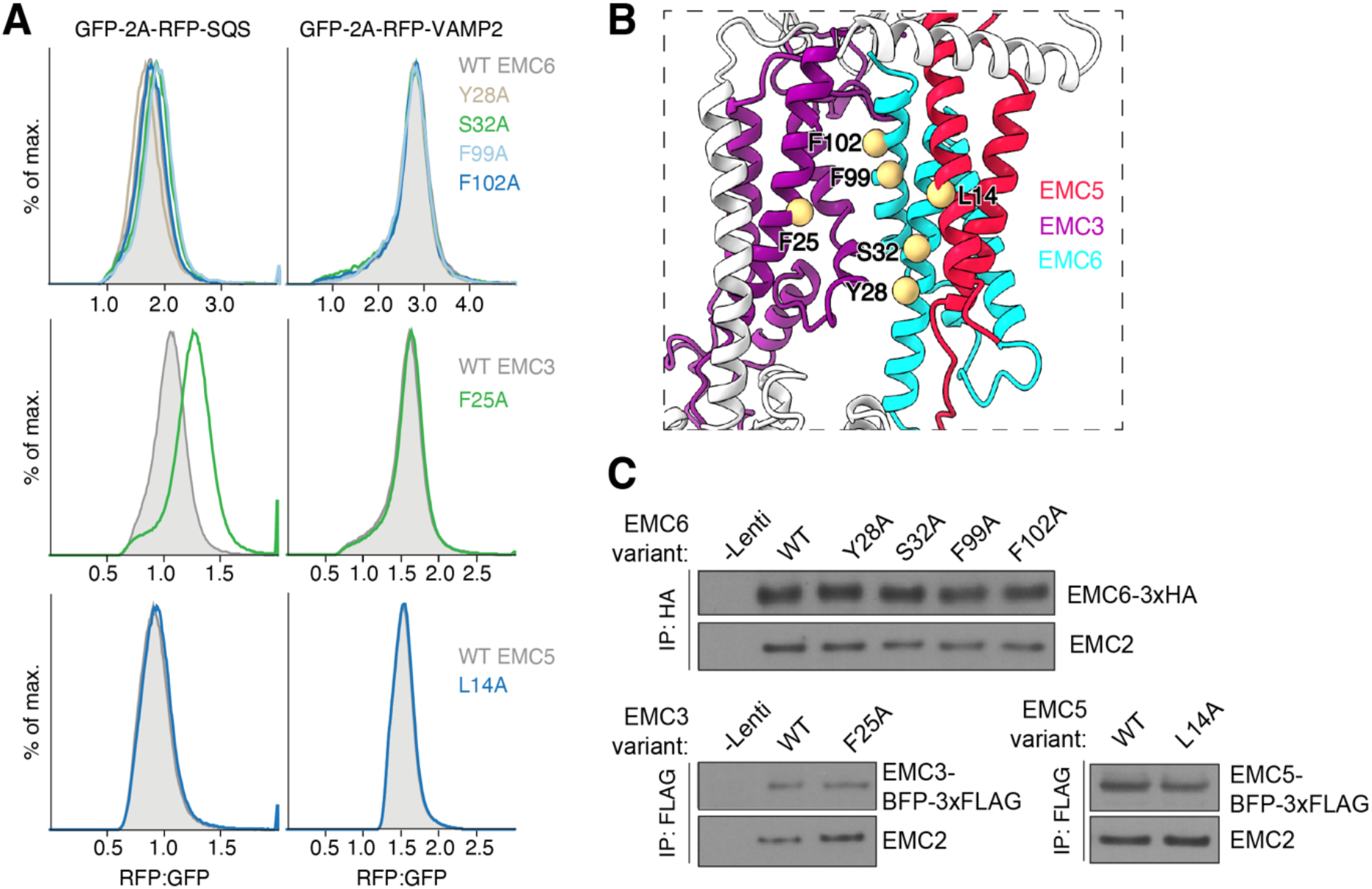
Mutation of the hydrophobic crevice does not impair SQS biogenesis in human cells. (**A**) HEK293 cells stably expressing RFP-SQS or -VAMP2 and cytosolic GFP as a normalization control were transduced with lentivirus to express the indicated mutants of EMC3, 5 and 6 in the hydrophobic crevice. The RFP:GFP ratio for each mutant was determined using flow cytometry and is plotted as a histogram. (**B**) Side-view of the membrane-spanning region of the EMC, focusing on the large cleft-like hydrophobic crevice. Residues on EMC3, 5 and 6 that were mutated in (A) line the cleft and are highlighted. (**C**) Incorporation of EMC subunit mutants into intact EMCs. A fraction of cells from (A) were harvested, solubilized, and subjected to anti-HA or anti-FLAG immunoprecipitation. Copurification with the soluble subunit EMC2 indicates successful incorporation of WT and mutant EMC3, 5 and 6 variants, suggesting that all of the mutant subunits are assembled into the mature EMC.

**Figure S3.**
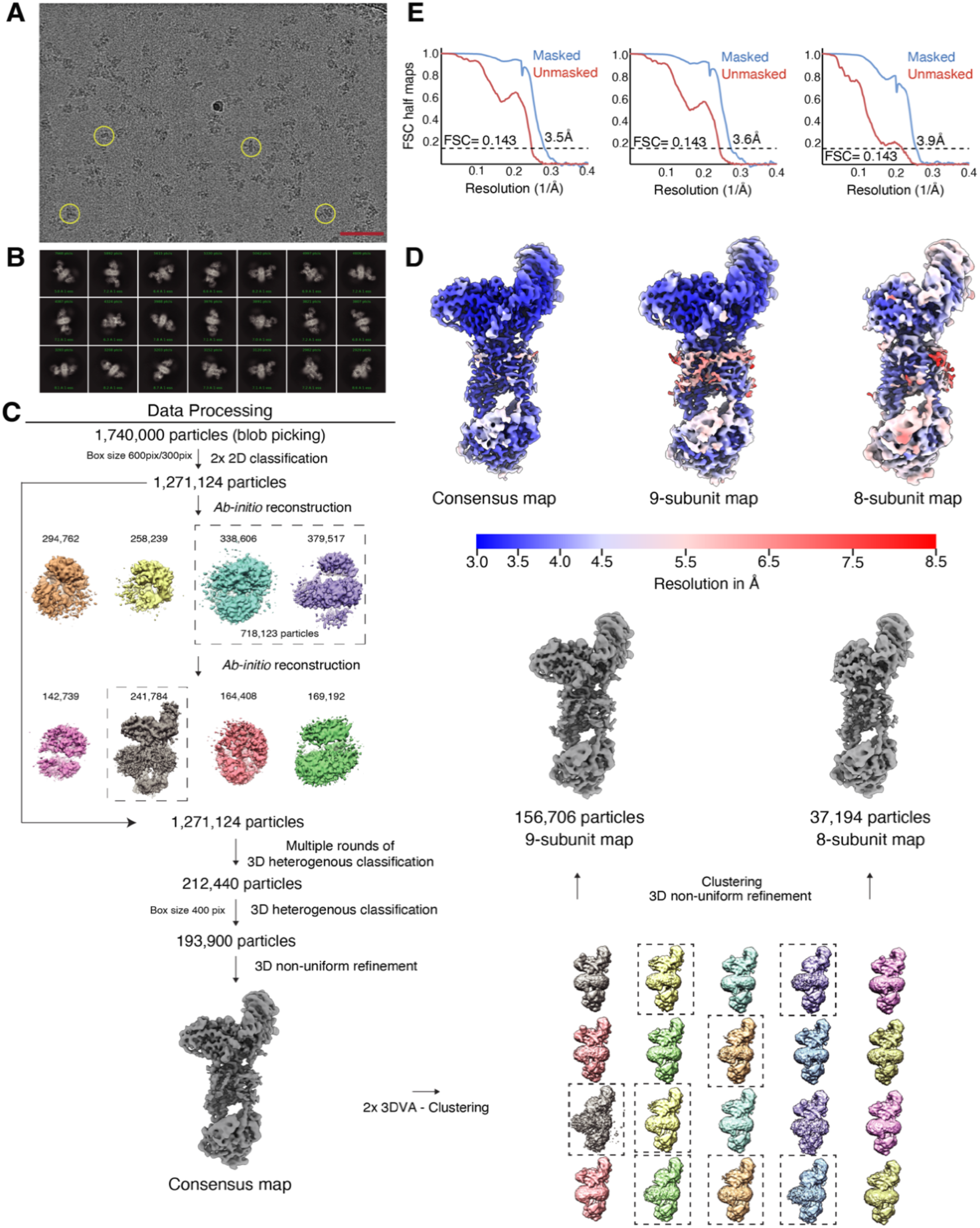
Classification and refinement procedure of an improved model of the human EMC. (**A**) A representative micrograph with several particles highlighted with yellow circles. Scale bar = 75 nm. (**B**) Representative 2D class averages generated during data processing. The number of particles for each class and its resolution are indicated. (**C**) Flowchart highlighting the data processing pipeline used to obtain an improved structure of the EMC. The 3D Variability Analysis (3DVA) enabled the exploration of the heterogeneity of the sample and allowed to parse out a subset of particles that lack the subunit EMC10, which provided unique insights into the placement of EMC10’s TMD. Particles with all nine subunits, or those missing EMC10 (dashed boxes) were combined separately. Particles with poorly-defined or low-resolution features were discarded (see Methods). (**D**) Final EM density maps colored by local resolution in Å. For clarity a dust filter was applied in ChimeraX. (**E**) Gold-standard Fourier Shell Correlation (FSC) curves for the consensus, 9- and 8-subunit complex maps generated by cryoSPARC V4.0.

**Figure S4.**
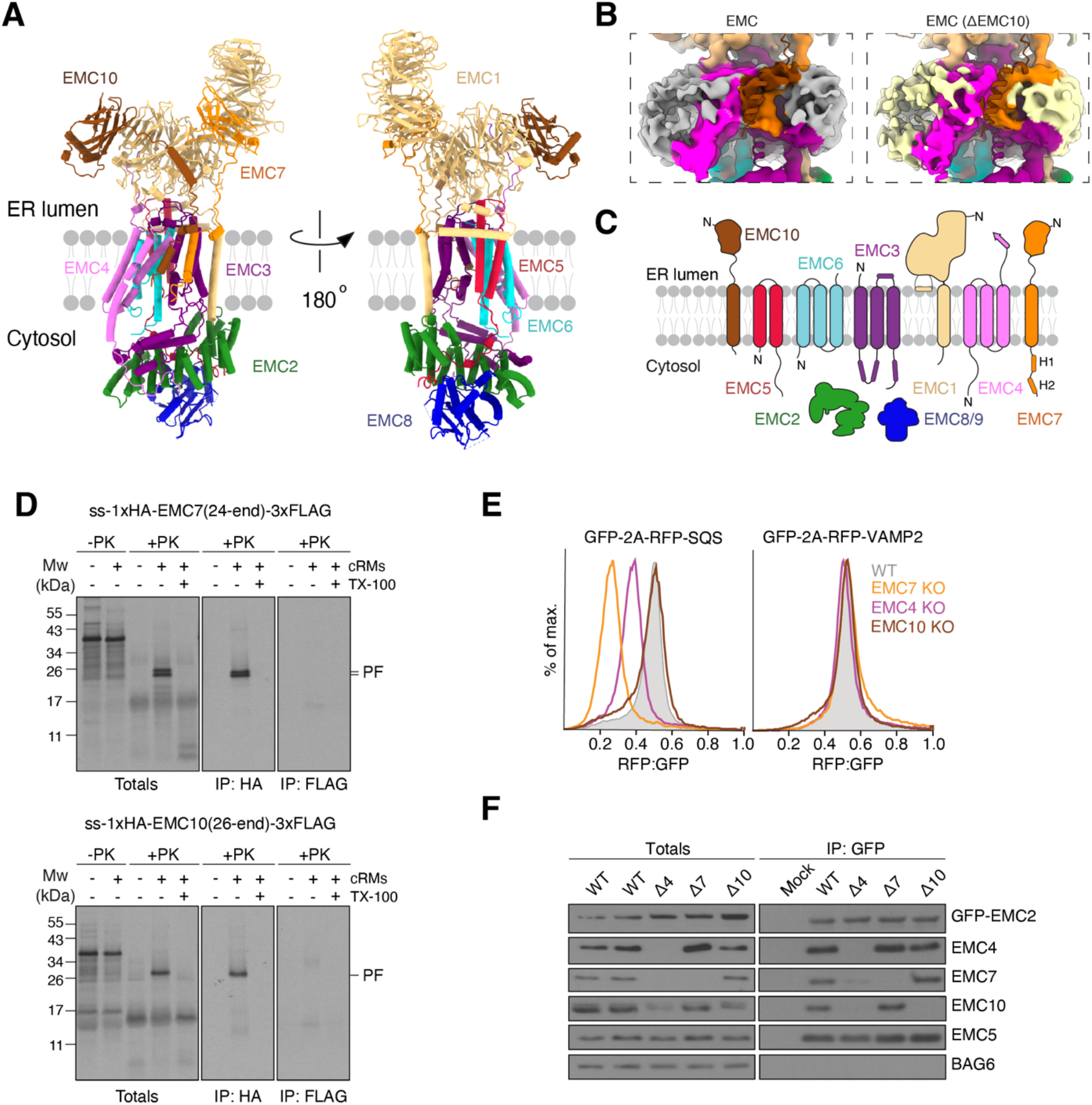
Architecture of the insertase-competent region of the EMC. (**A**) Updated model of the EMC, with views of the hydrophilic vestibule (left) and hydrophobic crevice side (right). (**B**) Low-pass filtered maps (5.5 Å) generated using volume tools in cryoSPARC V4.0. (Left) 9-subunit EMC complex map colored by the EMC subunits with the atomic model displayed as a superimposed cartoon. The EM density for the detergent micelle is displayed in gray. (Right) 8-subunit EMC complex (ΔEMC10) map. Due to the inherently flexible nature of EMC10’s TMD we could not unambiguously model its TMD, however, comparing +/Δ EMC10 maps gave insights into localization of its TMD because the ΔEMC10 map lacks additional density (colored in brown) enclosing the hydrophilic vestibule of the EMC. (**C**) Updated schematic of the topology of all nine EMC subunits. EMC8 and 9 are mutually exclusive paralogs. (**D**) EMC7 and EMC10 span the membrane. ^35^S-methionine labeled EMC7 (top) or EMC10 (bottom) carrying an N-terminal signal sequence (ss) and 1xHA tag, as well as a C-terminal 3xFLAG tag were *in vitro* translated in rabbit reticulocyte lysate supplemented with canine pancreatic rough microsomes (cRMs). Nascent chains were released from the ribosome with puromycin, and non-incorporated as well as cytosolically accessible proteins were digested with proteinase K (PK) in the presence or absence of Triton-X-100 to solubilize the cRM membrane. The resulting protease protected fragments were subjected to denaturing anti-HA and anti-FLAG immunoprecipitations (IP). Note that only the N-terminal HA tags of EMC7 and EMC10 were protected (PF = protected fragment) from PK digestion, whereas the C-terminal 3xFLAG was PK-accessible, indicating a type I, single-spanning topology for both subunits. (**E**) EMC4 and EMC7, but not EMC10 are required for SQS biogenesis in human cells. WT or EMC4/7/10 KO HEK293 cells were transduced with lentivirus to express RFP-SQS or -VAMP2. The relative level of the RFP-fused TA to an internal GFP expression control was measured via flow cytometry and plotted as a histogram. (**F**) Purification of EMC complexes from HEK293 cells stably expressing GFP-EMC2 (WT), with or without additional knockout of EMC4, 7 or 10. Samples of total lysate and elution following an IP via GFP-EMC2 were analyzed by SDS-PAGE and western blotting with the indicated antibodies.

**Figure S5.**
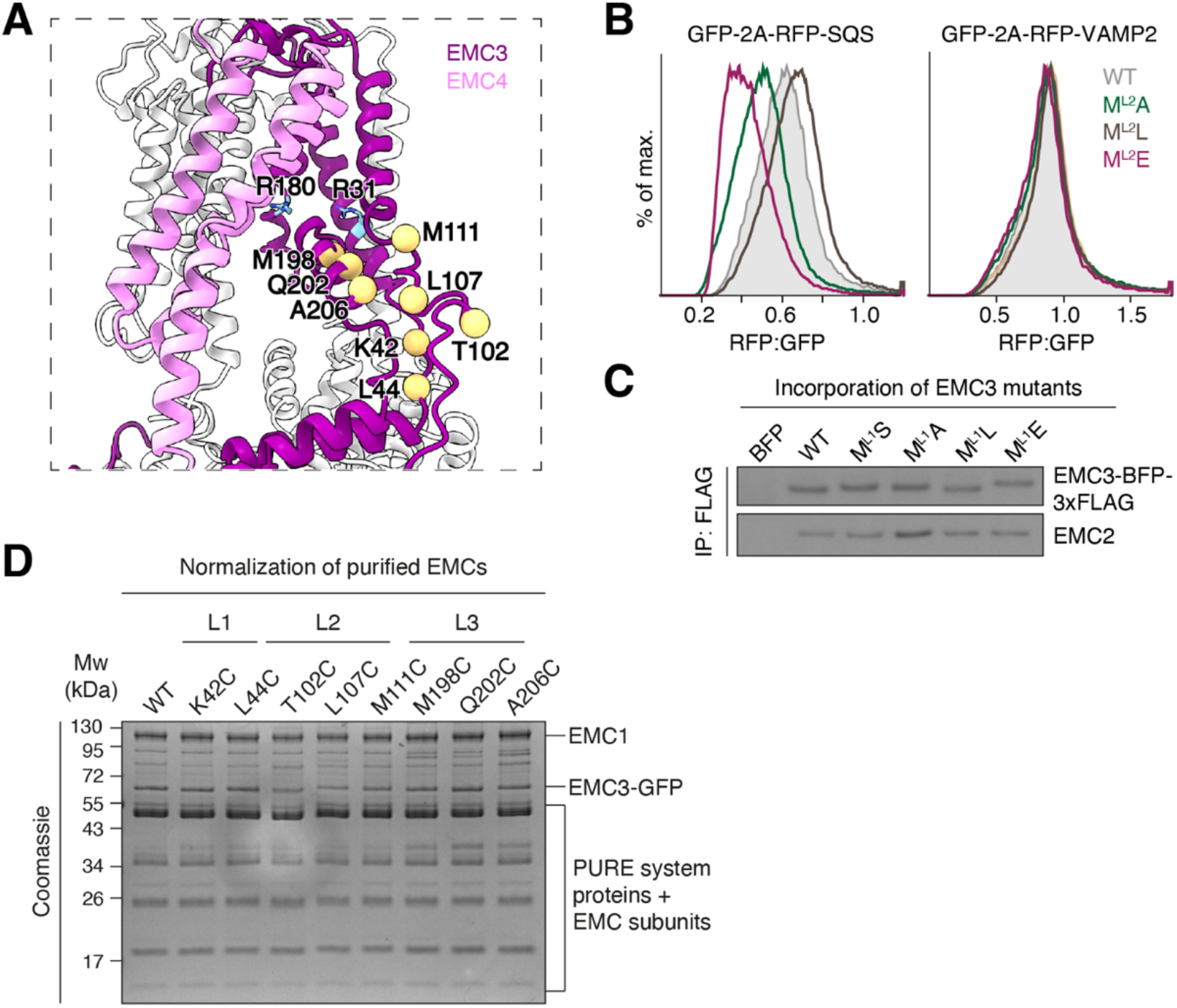
Substrate capture by EMC3’s hydrophobic loop 2. (**A**) View of the hydrophilic vestibule of the EMC, focusing on the three cytosolic loops of EMC3. Residues chosen to place cysteines for disulfide crosslinking are highlighted. R31 and R180 (in blue) are shown for orientation and EMC4, 7 and 10 are omitted for clarity. (**B**) HEK293 cells stably expressing RFP-SQS or -VAMP2 and cytosolic GFP as a normalization control were transduced with lentivirus to express the indicated EMC3 loop 2 mutants, along with BFP as a transduction marker. For each mutant, the RFP:GFP ratio of BFP-positive cells was derived via flow cytometry and is plotted as a histogram. M^L2^ refers to all four methionines in loop 2. (**C**) The indicated EMC3 loop 2 mutants were introduced into HEK293 cells via lentiviral transduction. Cells were harvested, solubilized and subjected to anti-FLAG immunoprecipitation (IP). Eluates were analyzed by SDS-PAGE and western blotting with the indicated antibodies. (**D**) Coomassie stained SDS-PAGE gel of the disulfide crosslinking experiment with purified EMC complexes shown in Figure 2B before analysis via autoradiography. The gel shows that equal amounts of EMC complexes were used in the different crosslinking reactions.

**Figure S6.**
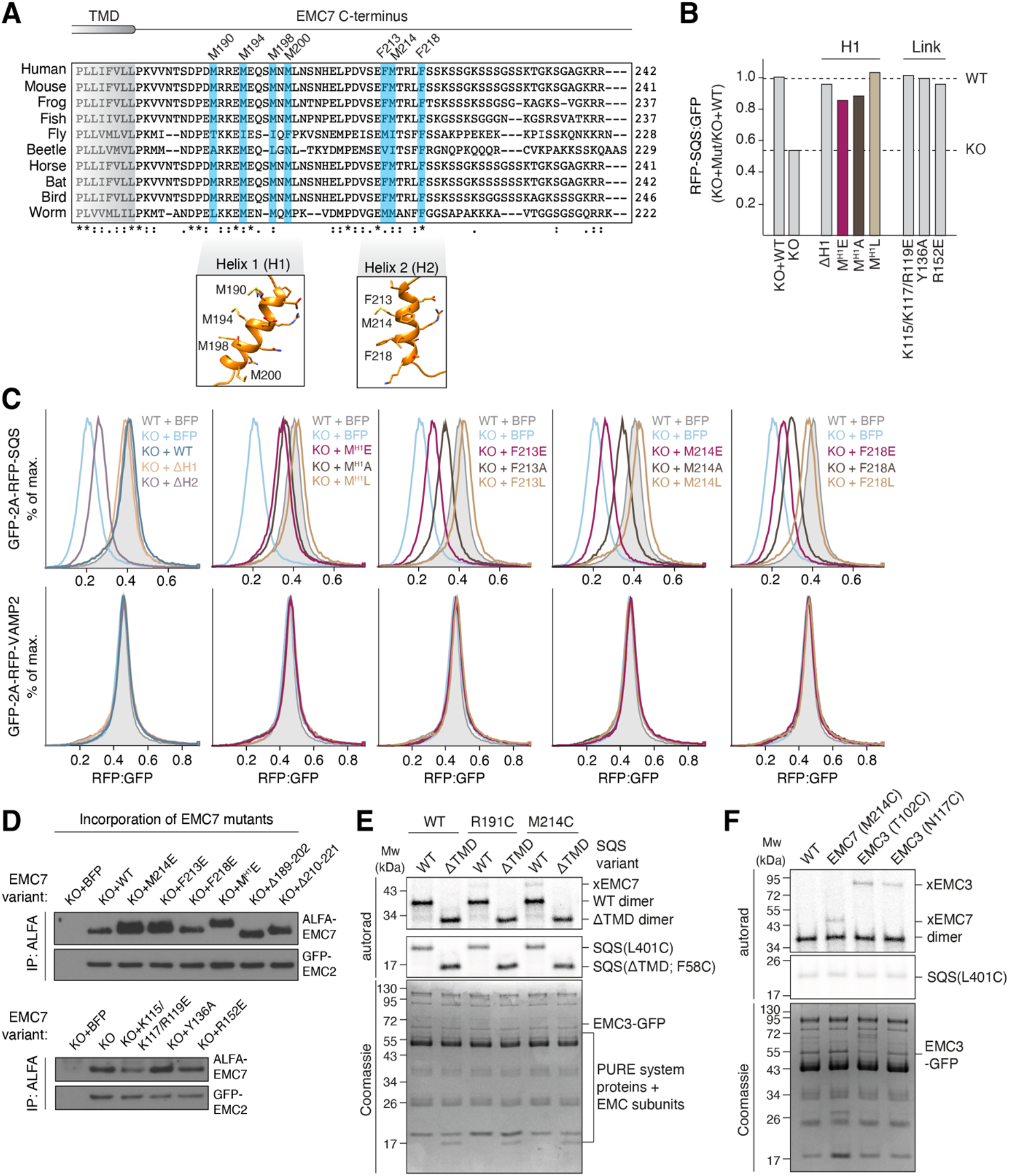
Substrate capture by EMC7’s cytosolic hydrophobic helix H2. (**A**) Alignment of EMC7 C-terminus sequences from various eukaryotes using Clustal Omega (Sievers et al., 2011). Two conserved sequence stretches are predicted by secondary structure algorithms to form α-helices, termed H1 and H2. Residues mutated in (C) are highlighted in blue. AlphaFold 2 models of EMC7’s H1 and H2 are shown below. H1 is methionine-rich and H2 is predicted to form an amphipathic α-helix. (**B**) As in Figure 2C, but with the indicated mutants of H1 or the lumenal linker (link) between the EMC7’s β-sandwich and TMD. M^H1^ refers to all four methionines in helix 1. (**C**) Wild type (WT) or EMC7 KO HEK293 cells were transduced with lentivirus to express either BFP alone or BFP plus EMC7(WT) or the indicated mutants. 48 h after rescue construct transduction, cells were transduced with lentivirus expressing either RFP-SQS or -VAMP2, as well as a cytosolic GFP normalization control. The RFP:GFP ratio was determined by flow cytometry and is plotted as a histogram. Note that deletion of H2 strongly impaired SQS insertion in cells. Mutation of hydrophobic residues F213, M214 and F218 on H2 to either alanine or glutamate, but not leucine, similarly impaired SQS, but not VAMP2 biogenesis. (**D**) A BFP control, wild type EMC (WT), or the indicated mutants of EMC7 were introduced into EMC7 KO HEK293 cells via lentiviral transduction. Cells were harvested, solubilized and subjected to anti-ALFA immunoprecipitation. Eluates were analyzed by SDS-PAGE and western blotting with antibodies against EMC2 and 7. (**E**) Purified EMC complexes containing either WT EMC7 or EMC7 with cysteines in H1 (R191C) or H2 (M214C) were incubated with purified CaM-SQS complexes with or without a TMD. The cysteine was placed either in the TMD (L401C) or the soluble linker (F58C), for the WT and ΔTMD SQS constructs, respectively. Disulfide crosslinking was carried out as in Figure 1E. (**F**) Disulfide crosslinking of SQS(L401C) with purified EMC complexes as above, containing cysteines either in H2 of EMC7 (M214S), loop 2 of EMC3 (T102C) or within the membrane (EMC3 N117C).

**Figure S7.**
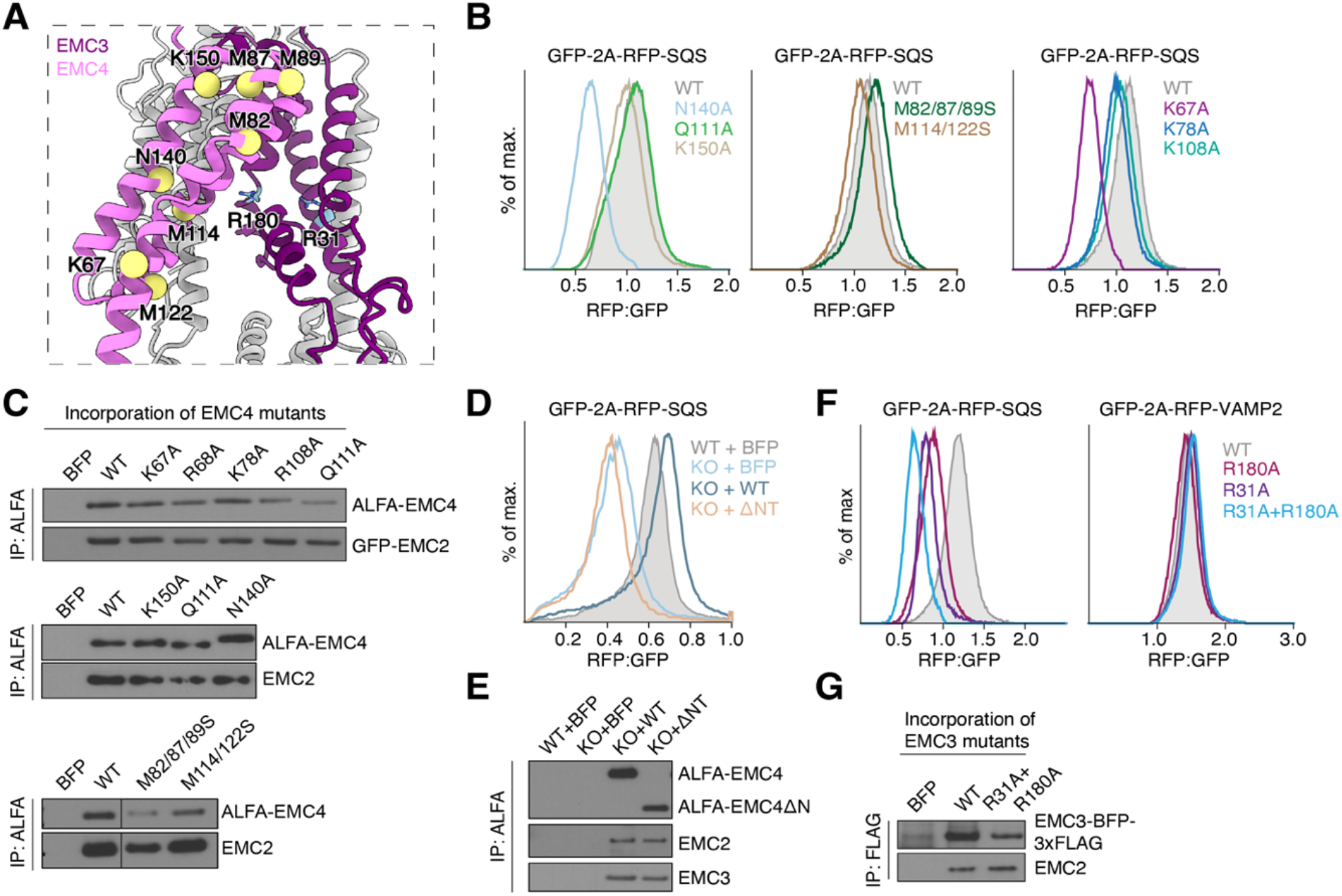
Biophysical properties of the hydrophilic vestibule. (**A**) View of the insertase-competent side of the EMC. EMC7 and 10 were omitted for clarity. Residues of EMC4 mutated in (B) are highlighted. R31 and R180 are shown as blue sticks for reference. (**B**) HEK293 cells stably expressing RFP-SQS and cytosolic GFP as a normalization control were transduced with the indicated mutants of EMC4, along with BFP as a transduction marker. The RFP:GFP ratio of BFP-positive cells for each mutant was derived via flow cytometry and is plotted as a histogram. (**C**) The indicated EMC4 mutants from Figure 2D and (B) were introduced into HEK293 cells via lentiviral transduction. Cells were harvested, solubilized and subjected to anti-ALFA immunoprecipitation (IP). Eluates were analyzed by SDS-PAGE and Western blotting with antibodies against EMC2 and 4. (**D**) The N-terminus of EMC4 is required for TA biogenesis in cells. HEK293 WT or EMC4 KO cells were transduced with lentivirus to express either BFP alone or BFP plus EMC4(WT) or a ΔNT mutant (residues 57-end). 48 h after rescue construct transduction, cells were transduced with lentivirus expressing RFP-SQS, as well as a cytosolic GFP normalization control. The RFP:GFP ratio of BFP-positive cells was derived via flow cytometry and is plotted as a histogram. (**E**) A portion of the cells from (D) was harvested, solubilized and subjected to purification of EMC4 variants via their N-terminal ALFA tag using the ALFA nanobody. The eluate was analyzed by SDS-PAGE and Western blotting with HRP-coupled ALFA nanobody or the indicated antibodies. (**F**) HEK293 cells stably expressing RFP-SQS or -VAMP2 and cytosolic GFP as a normalization control were transduced with lentivirus to express the indicated mutants of EMC3, as well as BFP. The RFP:GFP ratio of BFP-positive cells for each mutant was derived via flow cytometry and is plotted as a histogram. (**G**) A portion of the cells from (F) was harvested, solubilized and subjected to purification of EMC3 variants via their C-terminal 3xFLAG tag. Incorporation of the single mutants was described before (Pleiner et al., 2020).

**Figure S8.**
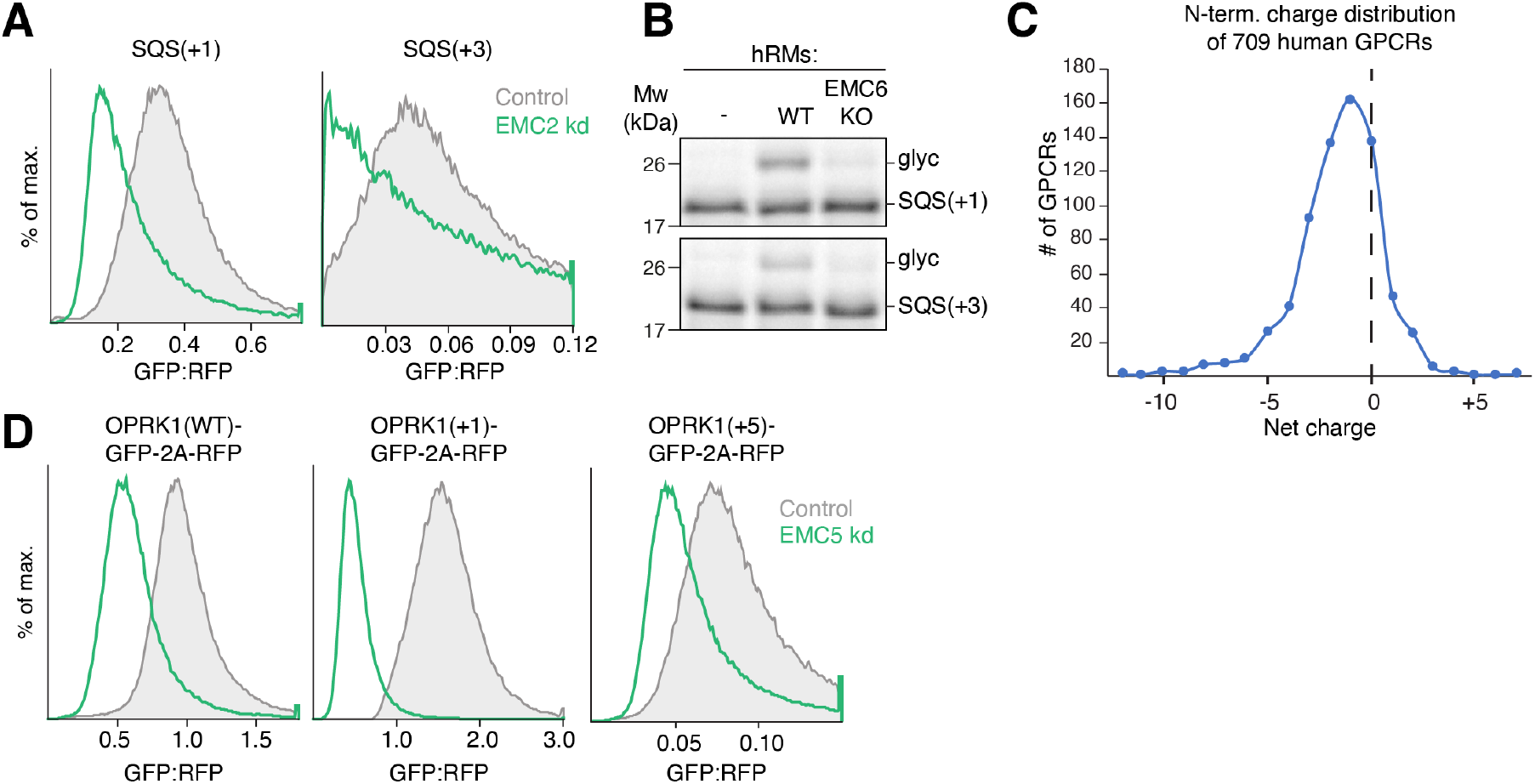
EMC dependence of SQS and OPRK1 charge mutants. (**A**) As in Figure 3D, but expressing the +1 and +3 C-terminal charge variants of SQS (as shown in Figure 3B). (**B**) Same assay as in Figure 3E, but with the +1 and +3 C-terminal charge variants of SQS (as shown in Figure 3B). (**C**) Distribution of N-terminal domain charge in 709 human GPCR sequences retrieved from the Uniprot database. Only GPCRs lacking a signal sequence (signal anchored) were chosen, since these are predicted to rely on the EMC for the successful translocation of their N-terminal domain and insertion of their first transmembrane domain in N^exo^ topology (N-terminus facing the ER lumen) (Chitwood et al., 2018). (**D**) WT (−5) or the indicated N-terminal domain (NTD) charge mutants of opioid receptor kappa 1 (OPRK1) were fused to GFP and expressed with RFP as a translation normalization marker in RPE1 cells. Cells were then treated with scrambled (control) or EMC5 siRNAs and analyzed by flow cytometry. Note that even though the stability of the positively charged NTD variants is reduced, they are still EMC dependent.

**Figure S9.**
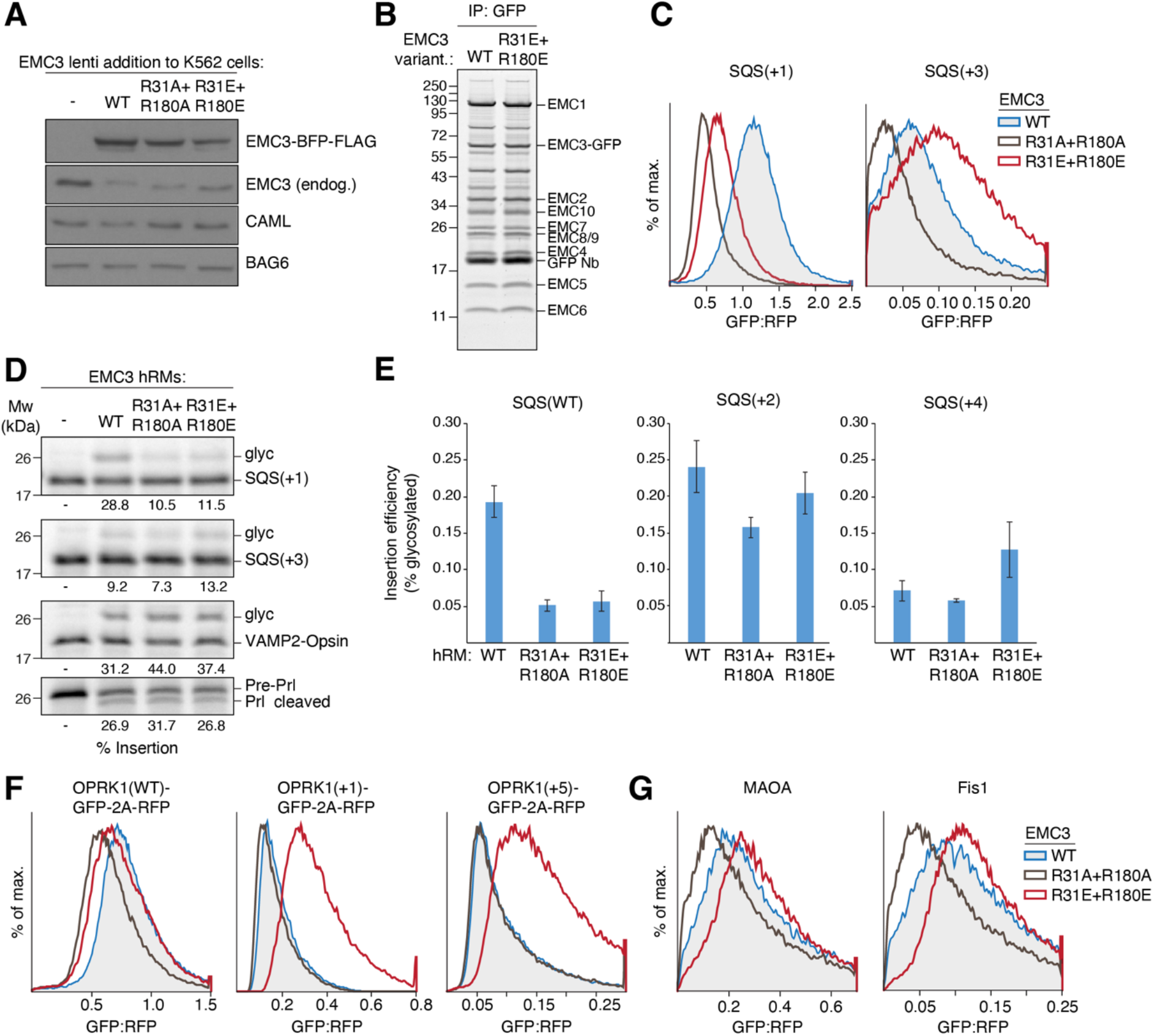
Charge reversal in the hydrophilic vestibule alleviates charge repulsion leading to misinsertion. (**A**) K562 CRISPRi ER GFP1-10 cells were transduced with lentivirus to express either EMC3 WT, R31A+R180A or R31E+R180E. Cells were harvested, solubilized and samples of the total lysates were analyzed by SDS-PAGE and Western blotting with the indicated antibodies. (**B**) Expi293 suspension cells stably expressing EMC3-GFP WT or R31E+R180E were solubilized and subjected to anti-GFP nanobody purification. The eluate was normalized by GFP fluorescence and analyzed by SDS-PAGE followed by Sypro Ruby staining. Note that both EMC3 WT and R31E+R180E mutant incorporate into EMCs with similar efficiency as they co-purify with all other EMC subunits. (**C**) Same assay as in Figure 4A, but expressing the +1 and +3 C-terminal charge mutants of SQS (as shown in Figure 3B). (**D**) Same assay as in Figure 4B, but with the +1 and +3 C-terminal charge mutants of SQS (as shown in Figure 3B). Expression of both EMC3 mutants does not impair the biogenesis of WRB-CAML dependent VAMP2 or the secreted protein prolactin (Prl) that depends on the Sec61complex (translocon). (**E**) Quantification of the experiment shown in Figure 4B. Error bars represent the mean and standard deviation from three independent replicates (n = 3). (**F**) WT (−5) or the indicated charge mutants of OPRK1 were fused to GFP and expressed with RFP as a translation normalization marker in RPE1 cells. Cells additionally expressed either BFP-tagged EMC3 WT, R31A+R180A or R31E+R180E. Cells were analyzed by flow cytometry to derive the GFP:RFP ratio of BFP positive cells. (**G**) Same assay as in Figure 4C, but expressing GFP11-tagged MAOA and Fis1.

